# Introducing selfisher: open source software for statistical analyses of fishing gear selectivity

**DOI:** 10.1101/2020.12.11.421362

**Authors:** Mollie E Brooks, Valentina Melli, Esther Savina, Juan Santos, Russell Millar, Finbarr Gerard O’Neill, Tiago Veiga-Malta, Ludvig Ahm Krag, Jordan Paul Feekings

**Affiliations:** National Institute of Aquatic Resources, Technical University of Denmark, Lyngby, 2800 Denmark; Thünen Institute of Baltic Sea Fisheries, Rostock, Germany; Department of Statistics, The University of Auckland, Private Bag 92019, Auckland, New Zealand

**Keywords:** catch comparison, covered codend, gillnet, mesh size, paired gear, trawl

## Abstract

Fishing gear is constantly being improved to select certain sizes and species while excluding others. Experiments are conducted to quantify the selectivity and the resulting data needs to be analyzed using specialized statistical methods in many cases. Here, we present a new estimation tool for analyzing this type of data: an R package named `selfisher`. It can be used for both active and passive gears, and can handle different trial designs. It allows fitting models containing multiple fixed effects (e.g. length, total catch weight, mesh size, water turbidity) and random effects (e.g. haul). A bootstrapping procedure is provided to account for between and within haul variability and overdispersion. We demonstrate its use via four case studies including (1) covered codend analyses of four gears, (2) a paired gear study with numerous potential covariates, (3) a catch comparison study of unpaired hauls of gillnets and (4) a catch comparison study of paired hauls using polynomials and splines. This free and open source software will make it easier to model fishing gear selectivity, teach the statistical methods, and make analyses more repeatable.

## Introduction

Fisheries aim to select for certain species and sizes of individuals while allowing others to avoid capture. Experiments are conducted to measure the selectivity of fishing gear and statistical models are used to characterize the selectivity patterns. The selectivity of fishing gear is commonly described by a retention curve, i.e. the probability of being retained in the net, which is usually a function of individual length or size and may vary between hauls (Wileman et al., 1996). Between-haul variation may be random, or it may depend on observed covariates such as total catch weight (Fryer, 1991; Suuronen & Millar, 1992; Erickson et al., 1996; O’Neill & Kynoch, 1996), environmental variables (He, 1993; Walsh & Hickey, 1993; Ryer & Barnett, 2006, Somerton et al., 2013), or the condition of individuals (Özbilgin et al., 2007; Ferro et al., 2008).

In some cases, catch data collected in selectivity studies could be analyzed with logistic regression methods, i.e. binomial generalized linear models, for which there are plenty of software options available. Binomial models are appropriate because, in many gear selectivity experiments, individuals end up in one of two compartments (e.g. codend vs cover; gear 1 vs gear 2; or test gear vs control gear; Wileman et al., 1996), i.e. there are two possible outcomes as in coin flips. However, a substantial amount of the analyses in this field are specialized and require specialized software. For example, obtaining confidence intervals on predictions typically requires accounting for extra-binomial variability (overdispersion) between and within hauls (Millar, 1993; Millar et al., 2004). Paired gear studies (where the test gear is tested against a control one with retention probability equal to one for the given species and lengths of interest) is another example that does not conform to a typical logistic regression model because the probability model is more complex as we will show below. This paper presents newly developed open source software that is specifically designed for modelling fishing gear selectivity, something that was previously limited. The package, `selfisher`, is implemented in the R statistical computing environment which is commonly used for many modern fisheries analyses (R core team 2020). The package is written with an interface that will be familiar to many users of regression methods in R. By making the software free and openly available, we aim to improve repeatability of analyses and enable teaching these analytical methods in classrooms for the next generation of fisheries scientists.

In this paper, first, we briefly describe the implementation of the `selfisher` R package. We describe how to install the latest version of the package and where to report bugs and where to reach out to other users with questions. Then, we provide a general description of how models are estimated by the package, including dealing with subsampled catches, a common occurrence in gear selectivity studies (e.g. Larsen et al., 2018; Melli et al., 2019; Veiga-Malta et al., 2020). We also address the common issue of overdispersion related to between-haul variability (Millar et al., 2004). We describe three general categories of statistical models, divided based on the mathematical probabilities underlying the estimation, while omitting details about experimental designs and code to run the models. Then, we describe the bootstrapping procedure used to account for variation within and between hauls in selectivity (Millar, 1993), potentially resampling from distinct pools of hauls based on gear type or tactic as in unpaired hauls (Herrmann et al., 2017; Savina et al., 2017). Then, we briefly describe four case studies used to illustrate the package’s capabilities, while providing more thorough descriptions with R code in online appendices. The case studies are (1) a covered codend study of haddock (*Melanogrammus aeglefinus*) with four different codends (O’Neill et al., 2016), (2) a paired gear study in a brown shrimp (*Crangon crangon*) trawl fishery where one trawl is nonselective (Santos et al., 2018), (3) a catch comparison study of unpaired hauls of gillnets avoiding an unwanted crab (*Cancer pagurus*) (Savina et al., 2017), and (4) a catch comparison study of paired hauls in a Norway lobster (*Nephrops norvegicus*) trawl fishery (Melli et al., 2018). Finally, we discuss current limitations of the `selfisher` package and potential future advancements.

## Implementation of selfisher

The `selfisher` package was designed to be flexible and robust for fitting and assessing a variety of gear selectivity models that can be represented with a binomial distribution. The code for `selfisher` was developed by modifying the R package `glmmTMB` (Brooks et al., 2017) because `glmmTMB` already had the capabilities needed for fitting and analyzing binomial mixed effect models. Previously, `glmmTMB` was developed by adapting the popular user interface from `lme4` (Bates et al., 2015) and increasing the model flexibility and fitting robustness by doing estimation with `TMB` (i.e. Template Model Builder). Prior to the development of `glmmTMB, TMB` was developed based on the algorithm of AD Model Builder (`ADMB`), which performs maximum likelihood estimation (MLE) in a fast and robust way (Fournier et al., 2012; Miller, 2013; Kristensen et al., 2016). The algorithm is fast and robust because it has information on the gradients of the likelihood surface via automatic differentiation. Additionally, `TMB` improves robustness by providing binomial and beta-binomial likelihood functions that are numerically stable even when probabilities are near zero or one. Thus, through inheritance, `selfisher` has a flexible user interface with `lme4`-style syntax and robust `TMB` code underlying the model estimation which is done using the same MLE algorithm as `ADMB`.

## Installing selfisher and where to ask questions

The package is continuously being improved and the most recent version can be found online in a GitHub repository. The current version of the package is 1.0.0. News about changes in each version can be found by typing news(package=“`selfisher`”)in an R console after installing the package. The package is mature enough that we do not expect to have changes that will affect existing models; most changes will be additions of new features (see Discussion below). The address of the GitHub repository is https://github.com/mebrooks/selfisher. There, you can find installation instructions and a forum for reporting bugs (i.e. issue tracker). Users are discouraged from posting questions on the issue tracker, which is reserved for bugs. For questions about `selfisher`, an email group is provided where users are encouraged to openly ask and answer each other’s questions (https://groups.google.com/d/forum/selfisher-users).

## Background of selectivity

All models in `selfisher` involve comparing the catches from two compartments (e.g. test vs control gear, gear 1 vs gear 2, or codend vs cover), which gives rise to data that can be analysed as a binomial response, subject to the use of appropriate methods to allow for overdispersion as described below. Due to technicalities of the underlying code, in `selfisher` syntax, the binomial response must be specified as a proportion (i.e. proportion of individuals of a given length in one compartment of one haul with respect to the total) and a total (i.e. total number of individuals of a given length in either compartment in one haul), as we will demonstrate in case studies.

There are three main categories of experimental designs in which each individual has two possible outcomes, i.e. studies that produce binomial data. The categories are (1) a selective net inside an outer nonselective small-mesh cover net (covered codend), (2) a selective net compared to a nonselective net (paired gear), and (3) a comparison between two selective gears (catch comparison). These can all be modeled using `selfisher` as we describe in sections below, but first we describe some generalities.

### Retention models

All three categories of analyses involve estimating a retention model; covered codend and paired gears experiments allow one to estimate the absolute retention, i.e. retention out of the population encountered by the gear, while in a catch comparison experiment the estimated retention is relative to that of a baseline gear. Regardless, the mathematical formulation is general. We use *r(I)* to refer to the retention model throughout this text, but as we describe in case studies below, it may depend on covariates other than length, *I*. See Table 1 for a description of all notations. As in binomial generalized linear models (GLMs), retention models use a link function to keep the retention probability in the range from zero to one. The most common link is the “`iogit`” (i.e. logistic), but other options include, “`probit`” (i.e. normal probability ogiv), “`cloglog`” (i.e. negative extreme value), “`loglog`” (i.e. extreme value/Gompertz), or “`Richards`”. The software default is the logit link. To fit retention model shapes that are more diverse than the built-in link functions, it is possible to use a logit link with more complex models such as polynomials (Holst & Revill, 2009) or smooth functions (Skalski & Perez-Comas,1993; Munro & Somerton, 2001; Fryer et al., 2003; Somerton et al., 2013) as demonstrated in case studies 3 and 4 below.

### Selectivity statistics I_50_ and SR

**Table 1.** Symbols

|  |  |
| --- | --- |
| $N_{i,h,l}$ | Number of fish caught in compartment $i$, haul $h$, in length class $l$ |
| $n_{i,h,l}$ | Number of fish sampled in compartment $i$, haul $h$, in length class $l$ |
| $s_{i,h,l}$ | Proportion of fish sampled in compartment $i$, haul $h$, in length class $l$ |
| $q_{h,l} = s_{i,h,l}/s_{j,h,l}$ | Subsampling ratio for compartment $i$ vs $j$ in haul $h$, in length class $l$ |
| $r(l)$ | Retention probability as a function of length $l$ |
| $p_h$ | Relative fishing power of test vs control gear in haul $h$ |
| $\phi_{paired,h,l}$ | Probability model in a paired gear model |
| $\phi_{compare,h,l}$ | Probability model in a catch comparison model |
| $\phi_{covered,h,l}$ | Probability model in a covered codend model |

Two estimated summary statistics of interest are the length with 50% probability of retention (I_50_) and the selection range (SR, i.e. the width of the range of length classes with 25% to 75% retention probabilities). Note that they only apply to models where retention probability monotonically increases with length, such as in covered codend and paired gear studies. In simple models with only length or size as a predictor of retention, then I_50_ and SR can be extracted from a model using the function `L50SR()`. In more complex models, such as the covered codend and paired gear case studies below, which involve additional covariates besides length, there are multiple ways to extract I_50_ and SR estimates. The covered codend case study demonstrates how to extract them algorithmically by finding the lengths that correspond to retention probabilities 0.25, 0.5, and 0.75 for each given value of covariates in the model. The paired gear example solves for I_50_ and SR mathematically using a model’s estimated coefficients. For either method, confidence intervals can be obtained by bootstrapping. In the future, a function will be added to `selfisher` which will be similar to the method demonstrated in the covered codend study.

### Subsampled catches

Often, in cases of abundant catches, it may not be feasible to measure the length of every individual that is caught and in those cases, only a fraction of the catch may be measured. This leads to additional statistical complexity in the analyses, but we have made `selfisher` capable of handling subsampling in any model. Here we denote the approximate fraction of individuals in compartment *i*, of haul *h*, and length class *I* that were sampled as *S_i,h,l_*. Although subsampling doesn’t always depend on *I*, we have written the package to be flexible enough to handle cases where each observed count (e.g. *n_i,h,l_*) has a different subsampling fraction. It is sufficient to include the ratio of subsampling fractions (i.e. the subsampling ratio) in a model, rather than each compartment’s fraction individually, *q^h,l^* = *S_i,h,l_/S_j,h,l_*, assuming here that *i* is the compartment being considered as “success” in the binomial context and *j* i s the alternative compartment. If raising factors were recorded in the data instead of subsampling fractions, then care should be taken when calculating the subsampling ratio to account for the fact that a raising factor is the inverse (1/*S_i,h,l_*) of a subsampling fraction. Subsampling fractions are between zero and one, while raising factors are greater than or equal to one.

The most general way to account for subsampling in a `selfisher` model is to specify the subsampling ratio *q_h,l_* using the argument `qratio` in a call to the `selfisher` function. In covered codend models with logit links (Millar, 1994) or catch comparison models with logit links (Holst & Revill, 2009) an `offset` could be used instead, but using `qratio` will be more broadly applicable because it can be used with any type of link and in paired gear models in addition to the other types.

## Three categories of experimental designs

### Covered codend

One way of characterizing the selectivity of towed gear is to capture the individuals that escape the net using a small-mesh cover, commonly known as the covered codend method. Then, the statistical model has strong information on retention because the total number of individuals encountered in each length class is directly observed. In this category, we compare the number of individuals sampled in the cover in haul *h* with length **I** (*n_1,h,l_*) and codend (*n_2,h,l_*) by modeling the proportion 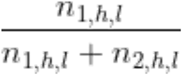 as the probability of being retained and sampled in the codend divided by the probability of being retained and sampled in the codend or escaping to the cover and being sampled (Millar, 1994): 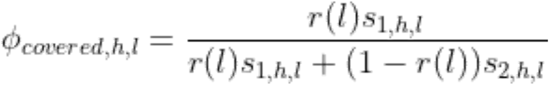 or more simply in terms of the subsampling ratio:

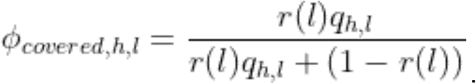

See Table 1 for definitions of all symbols.

### Paired gear (where one gear is nonselective)

Another way to characterize the selectivity of a fishing gear is to acquire knowledge on the population available to be caught in each haul. Paired gear studies accomplish this by deploying a control gear or codend (besides the one whose selectivity is being measured) that has full retention for the species and lengths of interest, i.e. all individuals of the given species entering that gear are retained. We assume that there is a probability, *p_h_*, of entering the test gear in haul *h*, given that the individual goes into either the test (*t*) or control (*c*) gear. The proportion of individuals observed in the test gear in haul *h* with length class *I* compared to the total number of individuals observed 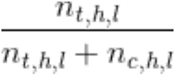 is modeled as the probability of entering the test net, being retained, and being counted, divided by the probability of entering either net, being retained and being counted:

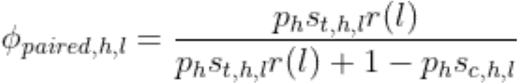

It’s convenient to write the model more simply in terms of the subsampling ratio:

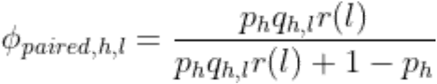

In paired gear studies, the ideal relative fishing power is 50% (i.e. p=0.5). If this is known a *priori* then it is possible to fix *p* at 0.5 by specifying `p formula=~0` i n the `selfisher` model. If another value of *p* is known a *priori* due to differences in effort, such as swept areas, then that can be specified using the `qratio` argument. For example, if the study has (haul-specific) subsampling fractions *st* and *sc* as well as (haul-specific) swept areas of *at* and *ac* for the test and control gear respectively, then one could use argument `qratio = at / ac * st / sc` together with `pformuia=~0` (e.g. Somerton et al., 2013).

### Catch comparison (where both gears are selective)

Catch comparison studies compare the catches in two gears, both of which are selective. Consequently, there is no direct information about the length distribution of the population being fished and it is only possible to model the relative retention probability given the population encountered during testing (Revill & Holst, 2004). In general, the relative retention probability model can be of arbitrary shape and, for example, may not be monotone.

The response variable is the proportion of fish retained by one gear versus the other

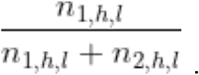

In a catch comparison study, the expected proportion of the total catch retained and sampled in gear 1 versus gear 2 is the probability of entering gear 1, being retained in gear 1, and being sampled over the probability of entering, being retained, and being sampled in either gear. It is always modeled with a logit link:

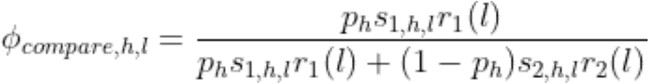

where *r_1_ (I)* and *r_2_(I)* are the absolute retention probabilities of the two gears, but it isn’t possible to estimate them separately. Holst & Revill (2009) showed that the expected proportion can be approximated with a polynomial with a logit link and an offset to account for any subsampling. They showed that the relative fishing pressure *p_h_* can be absorbed by the intercept and that it may vary randomly between hauls. The `selfisher` package allows fitting a random intercept of haul as in a mixed effects model. In general, the relative retention model in a catch comparison analysis can be formulated as either a polynomial (Holst & Revill, 2009) or a spline (Miller, 2013); see the catch comparison case studies in Appendices 3 and 4 for a demonstration of how this can be done with `selfisher`.

## Handling extra variability

### Overdispersion

Overdispersion is the presence of variability in the proportions that is in excess of the variability specified under the binomial model. Overdispersion can arise due to between-haul variability whereby the retention model varies from haul to haul. To a lesser extent it can also arise due to within-haul variability due to schooling behavior that violates the assumption that all fish behave independently. The accepted approaches for including this variability are the use of bootstrapping (Millar, 1993) and the use of mixed effects models (Millar et al., 2004). Both of these methods are available in `selfisher`. See the covered codend and paired gear case studies (Appendices 1 and 2) for examples of using mixed effects in `selfisher`. In this text, we do not get into the details of random effects because it is a large topic; however, note that in `selfisher`, they are implemented in the same way as `glmmTMB` (Brooks et al., 2017) using the Laplace approximation, which is a standard method commonly used in modern mixed modelling. All of the case studies demonstrate how to bootstrap as described below.

### Quantify uncertainty by bootstrapping

For any statistical model, it is important to compare predictions from the model - and the uncertainty around those predictions - with observed data to ensure that the model reasonably represents the data. A bootstrapping procedure was developed by Millar (1993) to account for variation between and within hauls and calculate appropriately wide confidence intervals. The bootstrapping method can account for overdispersion in data due to variability among hauls (as described above in the *Overdispersion* section); because of this, it is not necessary to include a random effect of haul in models to be bootstrapped. The bootstrapping method is also sometimes referred to as a “double bootstrap” in fishing gear selectivity literature, but this term has another meaning in statistics (e.g. Kuk, 1989). This method first resamples the same number of hauls from the observed set, with replacement. Then for each resampled haul, the method resamples observed fish within the haul. Then it refits the model to the resampled data set and the refit model is used to produce values such as predictions or parameter estimates. It is typically repeated one thousand times or more. This bootstrapping method is implemented in `selfisher` in a way that maintains all variables associated with each observed data point, not just length class, e.g. sampling fractions or total catch. This is facilitated by specifying the `haul` argument in the `selfisher` model fitting function; followed by a call to the `bootsel` function.

It is also possible to do resampling from pools of hauls, so that every bootstrapped dataset has the same number of hauls in each pool as in the original data (Herrmann et al., 2017). That is, if the original dataset has H_A_ hauls of type A and H_B_ hauls of type B, then it is possible to bootstrap in a way such that simulated data sets have H_A_ hauls of type A and H_B_ hauls of type B. This is done using the `pool` argument to the `selfisher` function. The inner part of the bootstrapping method (resampling fish within hauls) is the same as in the regular bootstrapping method. This is useful when hauls of the two gears or tactics being compared are unpaired. See the gillnet case study in Appendix 3 for an example of specifying pools of hauls.

### Bootstrapping and mixed modeling

Mixed modelling is a formal method that takes into account possible sources of variability in the data such as variation between hauls, enabling sound hypothesis testing and model selection. However, fitting mixed models can be computationally intensive. Moreover, the researcher is typically interested in obtaining overall selectivity predictions, rather than at the haul level, because these are relevant to the selectivity applied to the fishery. In that case it is necessary to refit the best candidate model, leaving out any random effects. Bootstrapping can then be used to obtain confidence intervals for estimated quantities such as predicted retention curves or I_50_ and SR. See case studies 1 and 2 for examples.

## Case studies

### 1. Covered codend analyses of four codends catching haddock

This case study uses the haddock data from an experiment that employed the covered codend method to investigate the selective performance of four codends (O’Neill et al., 2016). The codends were made from netting materials with different twine bending stiffnesses and mesh sizes.

We begin the case study by looking at just one gear type to demonstrate different link functions that can be used and to show how to account for subsampling, which in this example varies with length. The default link is the logistic, but we also consider the probit and Richards curves and a spline. Having chosen a model, we bootstrap to estimate 95% confidence intervals for the proportion retained by the codend.

We then analyse all four gear types together and investigate the influence of length, mesh size, bending stiffness and catch size. We assume the principle of geometric similarity (as used by Tokai et al. (1996) to investigate grid selection) and explore a number of models and choose the best fit using Bayesian information criterion (BIC). When choosing the best model, we include a random effect of haul to account for between-haul variation. As in the original publication, we show that selection is dependent on all three parameters. Before bootstrapping, we drop the random effect of haul from the best model because the bootstrapping method accounts for between-haul variability and the random effect would slow it down considerably. We bootstrap to estimate confidence intervals for the proportion retained by each gear and numerically solve for I_50_ and SR dependent on covariates. See Appendix 1 for details and code.

### 2. Paired gear analyses of codend selectivity dependent on mesh size

This case study draws on a subset of data from the German research project CRANNET (Santos et al., 2018). The experimental method consisted of fishing with two identical beam trawls, simultaneously and in parallel on the same shrimp population. One of the trawls mounted a small-mesh (11 mm) control codend with very limited selectivity, assumed to be nonselective on the range of shrimp lengths available for the trawl. The second trawl mounted a test codend. The subset of data analyzed here consists of catch data from 87 hauls, during which 13 diamond-mesh codends varying in mesh size ranging from 19.1 mm to 36.3 mm were tested. The goal was to model I_50_ and SR as a function of mesh size, and to quantify any effect of two additional haul covariates, sea state and catch weight.

The statistical modeling of selectivity begins with a mixed model to formally assess the effect of mesh size, sea state and catch weight, while controlling for random variation among hauls. *A priori*, the assumption of geometric similarity (that is, I_50_ and SR being proportional to mesh size) was assumed to be the default model, and it is shown that this corresponds to using a `I (length/meshsize)` term in the `selfisher` formula interface for the retention model (Baranov, 1948). The default model was compared to several others and found to be preferred (using BIC), and neither sea state nor catch weight had a significant effect.

Having chosen geometric similarity (with respect to mesh size) as the preferred model, this model was refitted without random effects so as to estimate size selectivity at the population level. Bootstrapping was used to obtain appropriate confidence intervals on I_50_ and SR for any given mesh size.

In addition, this case study demonstrates the use of `psplit=TRUE` (unequal fishing power of the paired codends), the use of sampling ratios, use of the `inits()` function to specify good starting values (because without it some models converged to local minima that didn’t make any sense), and shows how I_50_ and SR can be obtained directly from the model fitted by `selfisher`. See Appendix 2 for details and code.

### 3. Catch comparison analyses of unpaired hauls of gillnets avoiding an unwanted crab

This example deals with data from an experiment originally published by Savina et al. (2017). Two soak tactics (12h at day and 12h at night) were compared in the Danish gillnet plaice fishery to estimate the role that the choice of a soak tactic plays in the catch efficiency of both target and unwanted species. This is a subset of the original dataset (one species, two soak tactics) where we are looking at the unwanted invertebrate edible crab (*Cancer pagurus*).

We use the method developed by Herrmann et al. (2017) which was developed for assessing the effect of changing the gear design on the relative length-dependent catch efficiency. This example is representative of experimental fishing where the catch data obtained for two gears or tactics were not collected in pairs, and can allow for a different number of deployments.

This case study is a typical model for catch comparison of multiple haul data without subsampling using a spline. To get confidence intervals on predictions, we bootstrapped from two pools according to tactic (Night vs Day) using the argument `pool=tactic` to the `bootsel` function. See Appendix 3 for details and code.

### 4. Catch comparison analyses of paired hauls of *Nephrops* twin-rigged trawls

The example is based on the data from Melli et al. (2018). An anterior gear modification, namely the counter-herding device FLEXSELECT, was tested in a twin-rig configuration, where two identical trawls were towed in parallel. One trawl was equipped with FLEXSELECT, referred to as the test trawl, while the other worked as baseline. The aim of the study was to determine if FLEXSELECT could reduce the fish bycatch in a *Nephrops-directed* fishery. The data used in the example are from haddock, which was found to be strongly affected by the counter-herding device.

Following the steps of the published paper, we conducted a catch comparison analysis, modeling the relative retention as a 4th-order polynomial. In addition, we used a spline with 4 degrees of freedom using the `splines` package and performed model selection to determine if it fitted the data better. Considering that part of the hauls were conducted in day-time and part in night-time, “time” was included in the model as an explanatory variable to determine if the length-based efficiency of FLEXSELECT presented diel differences. We predicted both catch comparison rates and catch ratio with bootstrapped confidence intervals using the `predict` and `bootsel` functions from `selfisher`. See Appendix 4 for details and code.

## Discussion

We have introduced an open source R package for estimating fishing gear selectivity of both towed and passive gear, making it easier for anyone to analyze fishing gear selectivity data without writing extensive amounts of code. We have demonstrated its broad applicability in four case studies spanning a range of experimental designs. The case studies have shown that results from `selfisher` are comparable to previously published results and that `selfisher` is more flexible than some methods (e.g. a single model to quantify the effect of changing mesh size on I_50_ and SR). Some of the features of `selfisher` that were demonstrated in the case studies are summarized in Table 2. The case studies aim to demonstrate best practices based on current knowledge. However, this is an active area of research and with a new powerful model fitting tool, best practices may change. Even with (or especially with) a powerful tool, analyses require careful thought and checking of results. For example, in complicated models such as paired gear models which contain two submodels (retention and relative fishing power), it may be necessary to be cautious about identifiability of parameters and local optima encountered during maximum likelihood estimation, but better starting values help avoid those issues as demonstrated in Appendix 2 (Bolker et al., 2013).

**Table 2.**
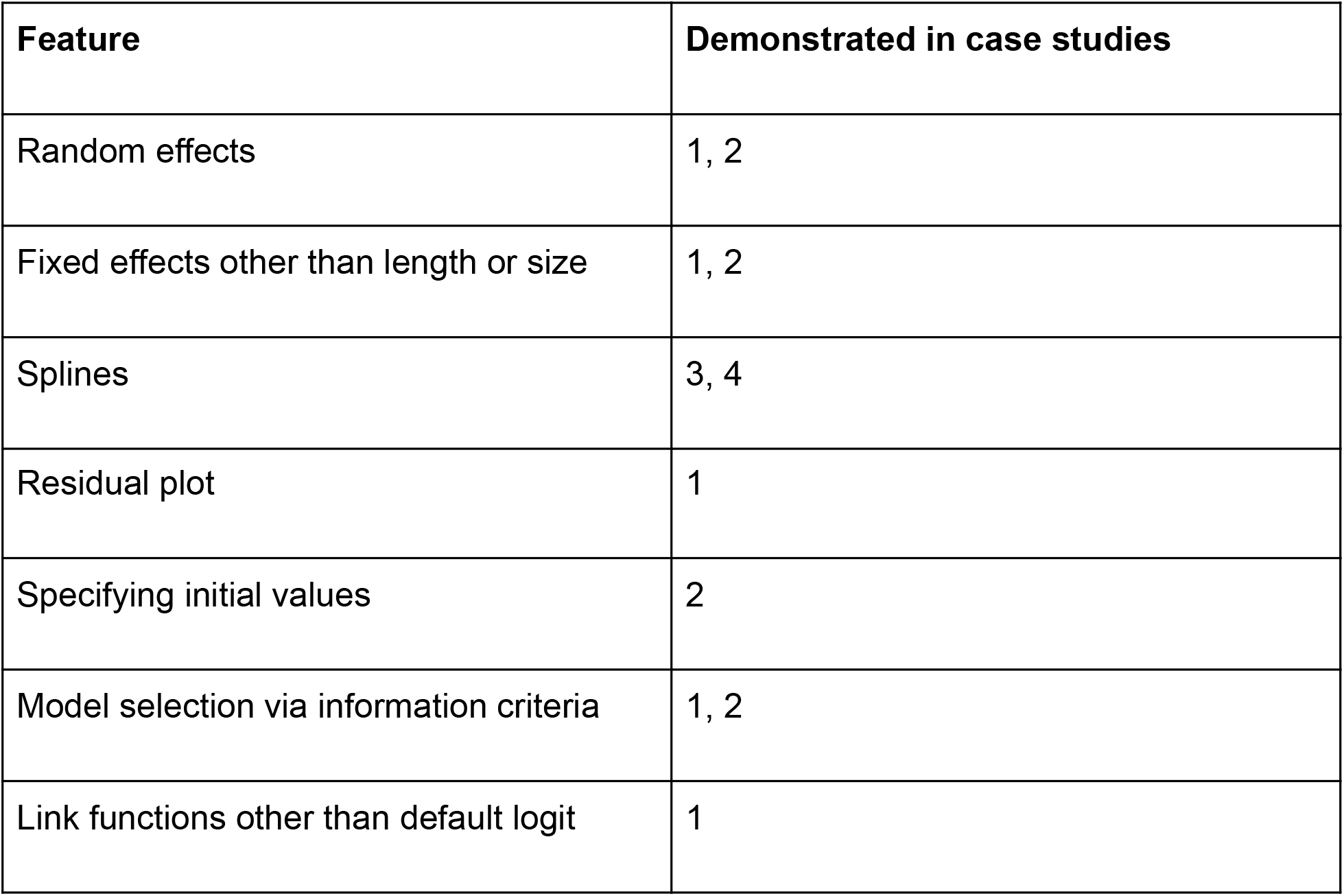
A list of `selfisher` features demonstrated in the four case studies with code in Appendices 1 through 4 (respectively).

We have several advancements for the package either planned for the future or already underway. We plan to add functions to calculate discard ratio indices and indicator functions (e.g. Wienbeck et al., 2014; Santos et al., 2016; Veiga-Malta et al., 2019). We may add a function to facilitate model averaging, although it is currently possible to piece this functionality together with the existing features. We have not tried to fit structured non-monotonic curves (e.g. bell-shaped curves of gillnet absolute selectivity based on geometric similarity, Baranov 1948) with `selfisher`, but we will explore this possibility in the future. We will investigate how to choose starting values of parameters in models that have Richards link, to increase robustness. We plan to implement a general method to extract I_50_ and SR from complex models as demonstrated in case studies 1 and 2. To handle overdispersion more elegantly, we plan to add the option of having a beta-binomial distribution for the response (Miller, 2013). We are already in the process of developing a Shiny app, which will facilitate simple standard analyses without the need for writing code; this will help bridge the gap for scientists or managers with extensive experience in gear development but little experience with R. As an open source package, code developers are encouraged to contribute improvements through GitHub such as those listed here.

Having access to a free and open source software should benefit this field of research in several ways. It allows researchers to share code and thereby foster a community for discussion and repeatability. The free nature of the software will enable researchers and managers with limited budgets - such as those in developing countries - to perform analyses themselves. It gives statistical methods of retention modelling a way into classrooms containing the next generation of fisheries scientists who are already learning modern regression methods as part of a general scientific curriculum.

## Acknowledgements

We thank William T Stockhausen for the trick of fixing *p_h_* to any number in a paired gear

analysis. We thank the helpful crews of both research and commercial vessels, as well as the fisheries technicians of the research institutes providing essential support during the fishing trips. We are grateful for funding from the European Maritime and Fisheries Fund (EMFF) and the Ministry of Environment and Food of Denmark (Miljø-og Fødevareministeriet) as part of the projects FastTrack - Sustainable, cost effective and responsive gear solutions under the landing obligation (33112-P-15-013), FastTrack II - Sustainable, cost effective and responsive gear solutions under the landing obligation (33112-P-18-051), and Flexselect - Innovative and Flexible solutions for the Nephrops Fisheries (33113-I-16-068).

## Data Availability Statement

Data used in the case studies is available as part of the `selfisher` package on GitHub. See the *Installation* section above as well as the appendices.

## Appendix 1 Covered codend analyses of four codends catching haddock

This example deals with data from an experiment published by O’Neill et al. (2016) that investigated the selectivity of haddock (*Melanogrammus aeglefinus*) in four codends made from netting materials with different twine bending stiffnesses and mesh sizes. Three of the codends had a nominal mesh size of 120mm and one a nominal mesh size of 130mm. The twine bending stiffness values were in the range 0.64 to 1.1kN *mm^2^*. We label the codends as 120low, 120med, 120high and 130med to reflect their mesh size and bending stiffness (as categorised by the netting manufacturers). As in the original analysis, we show that selection is dependent on both of these parameters and the total codend catch weight.

### Preliminaries

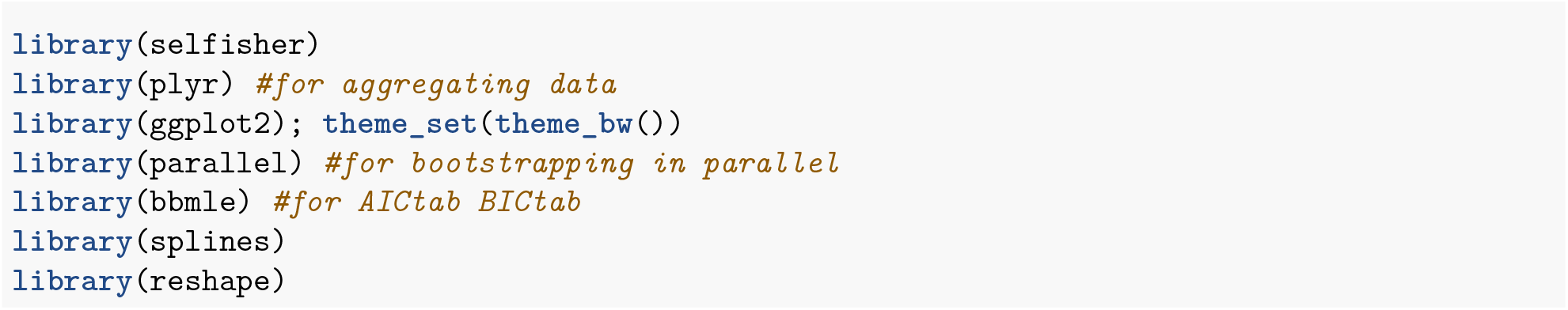

### Data structure

We load the data each row of which corresponds to the fish of a given length from a given haul. The length (cm), haul number, mesh size (mm), twine bending stiffness (*kNmm^2^*), number of fish measured from the codend, codend raising factor, number of fish measured from the cover, cover raising factor, catch size (kg) and codend label are specified respectively.

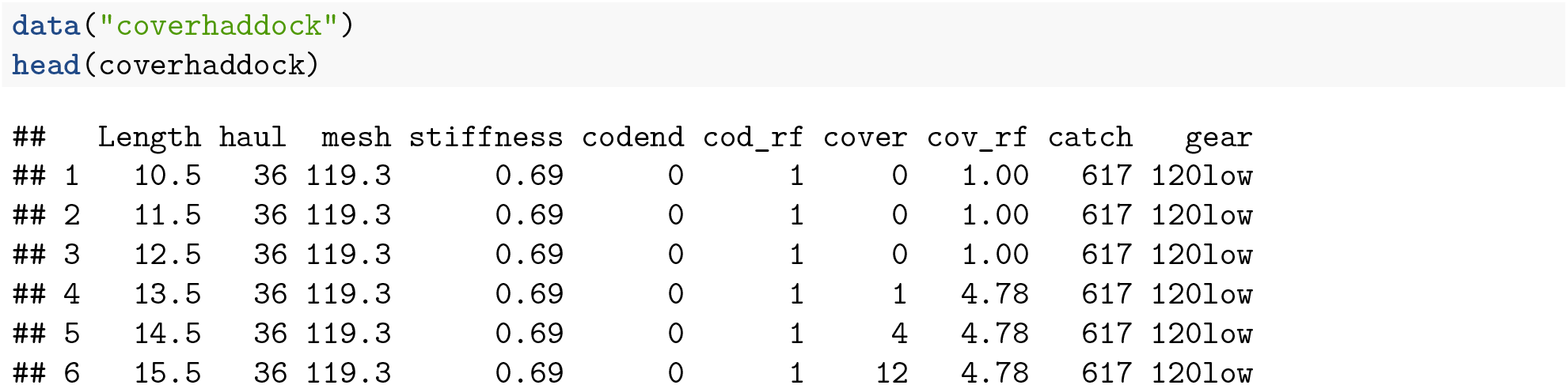

Here we can see that the raising factor varied by length class, which is not a problem in `selfisher`.

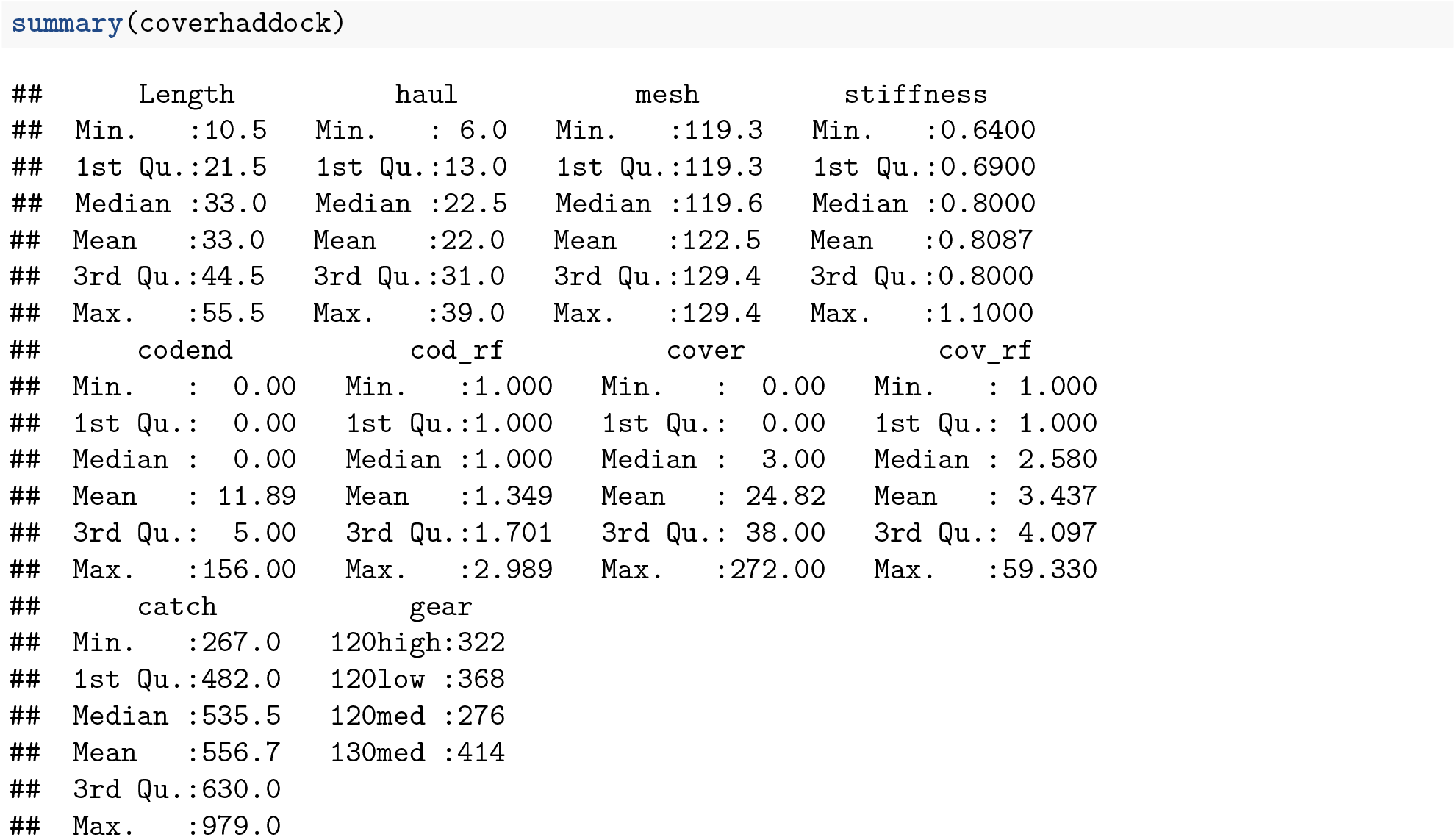

We can also see that all the hauls are contained in one data frame. The data is organized into what is called “long format”.

### Transforming data

For a model in `selfisher`, we need to convert counts into proportions and totals. Unlike other GLM functions for binomial regression, it is **not** possible to specify the binomial variable as a two-column response variable, e.g. `cbind(N_test, N_cover)`.

Because not all fish in the samples were counted, we will account for this in the model (Millar 1994). The values we need in the model are calculated as `cov_rf/cod_rf`. If instead we had a sampling fraction, we would calculate `qratio = sampling_test/sampling_cover` because a sampling fraction is the inverse of a raising factor. We create a new column in the data with the value of `qratio = cov_rf/cod_rf` for each row. An easy way to compute this value by row is to use the `transform` function.

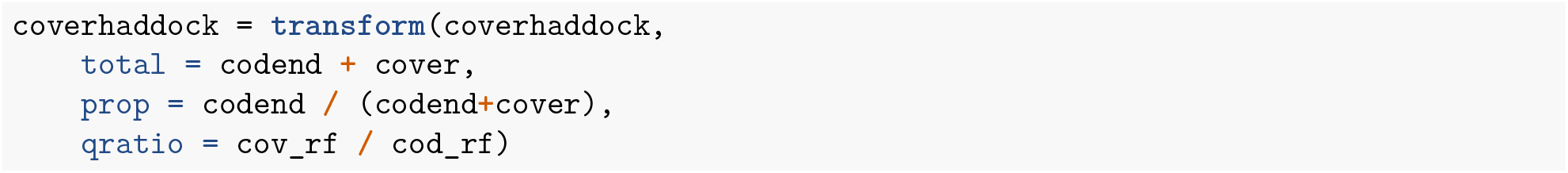

We drop rows of data where no fish were observed because they don’t contain any information (i.e. where total = 0). This doesn’t affect the model except to allow for bootstrapping later.

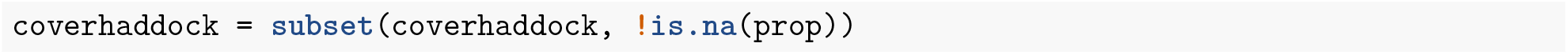

### Single gear model with covered codend

We will start out with a simple example of one gear type (“120low”). So we need to subset the data.

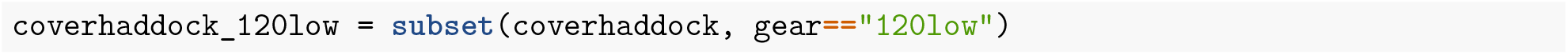

The following is a model for multiple haul data from a covered codend experiment with subsampling. It could be argued that there should be a random effect of haul in the models to account for variation among hauls and avoid pseudoreplication, but we will keep it simple for this first example and only include a fixed effect of length.

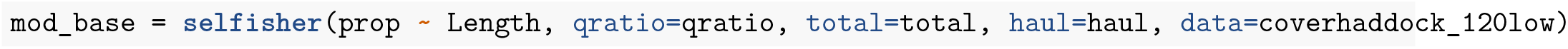

### Extracting residuals and other standard methods

We can check the model residuals for patterns. There’s a residuals function for `selfisher` models. All methods for `selfisher` models can be displayed as follows:

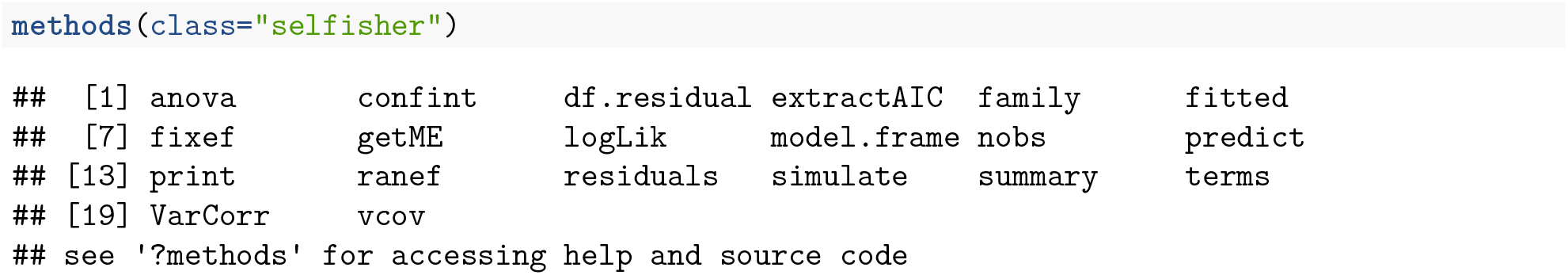

Here is one way to plot the residuals.

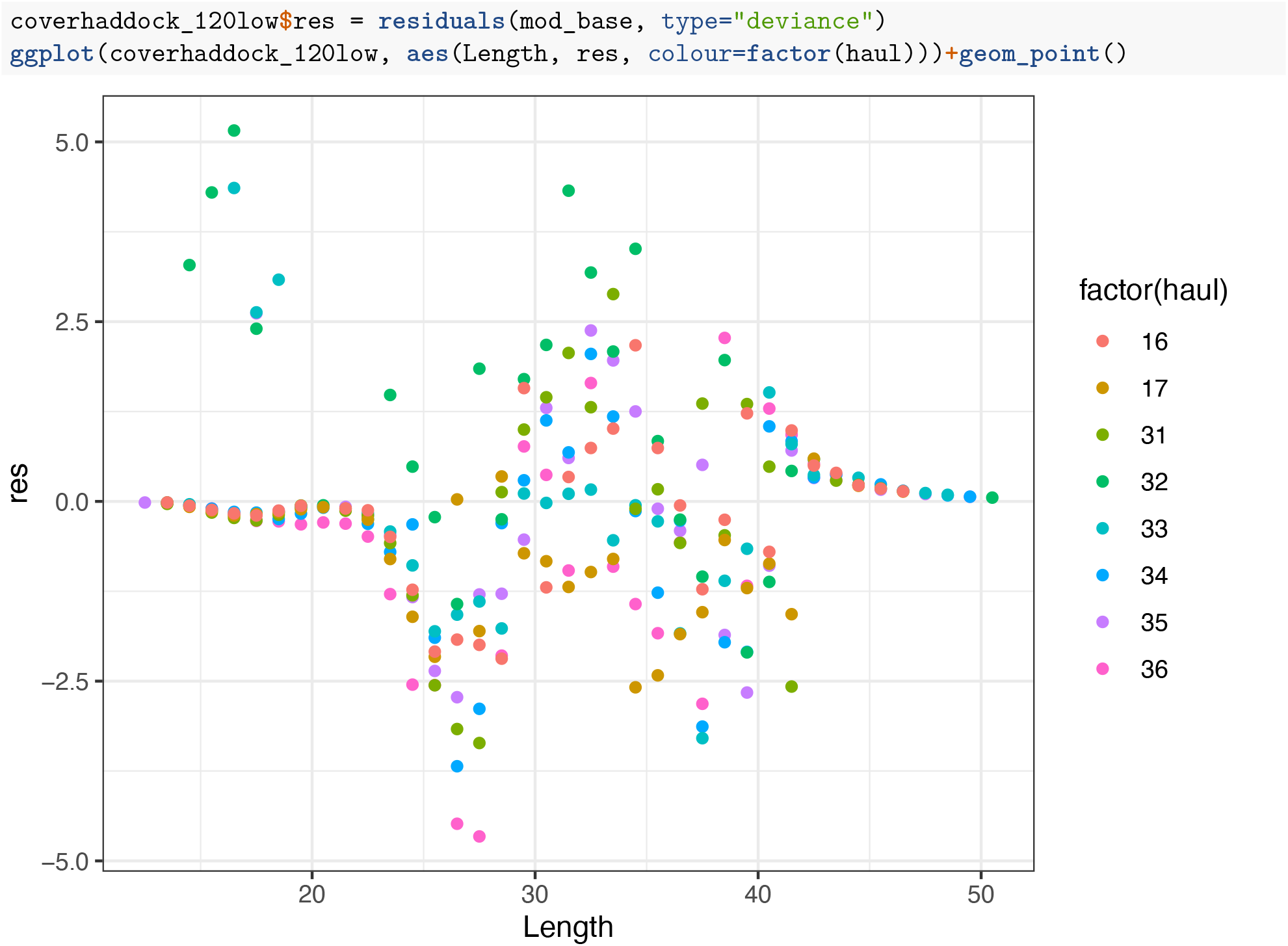

Residuals from GLMs are notoriously opaque, but in the future, we will try to make `selfisher` compatible with the `DHARMa` package to make it easier to assess residuals (Hartig 2020).

### Links other than logit

It is also possible to consider other link functions, or use a spline. See the function documentation (`?selfisher`) for a list of implemented link functions. Here we fit the logistic, probit and Richard’s curve and a spline.

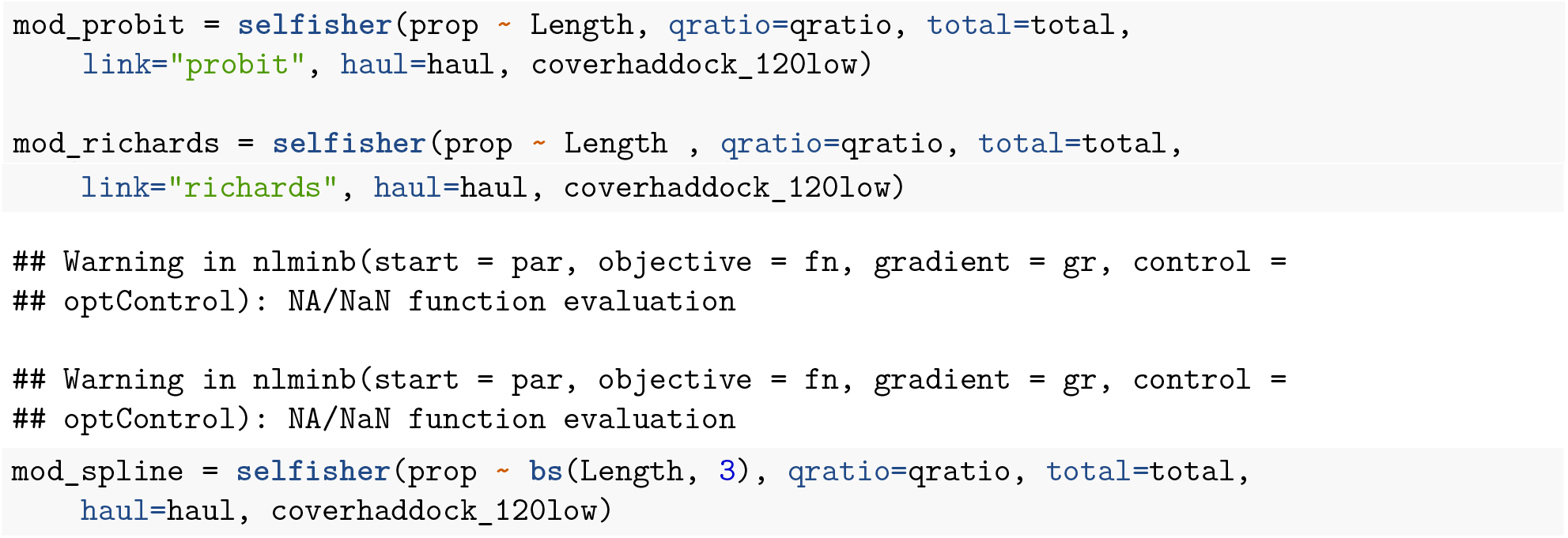

Fitting the model with `link=“richards”` produced some warnings, but this is ok. The model is valid if the `summary` function is able to produce non-NA standard-errors as seen below.

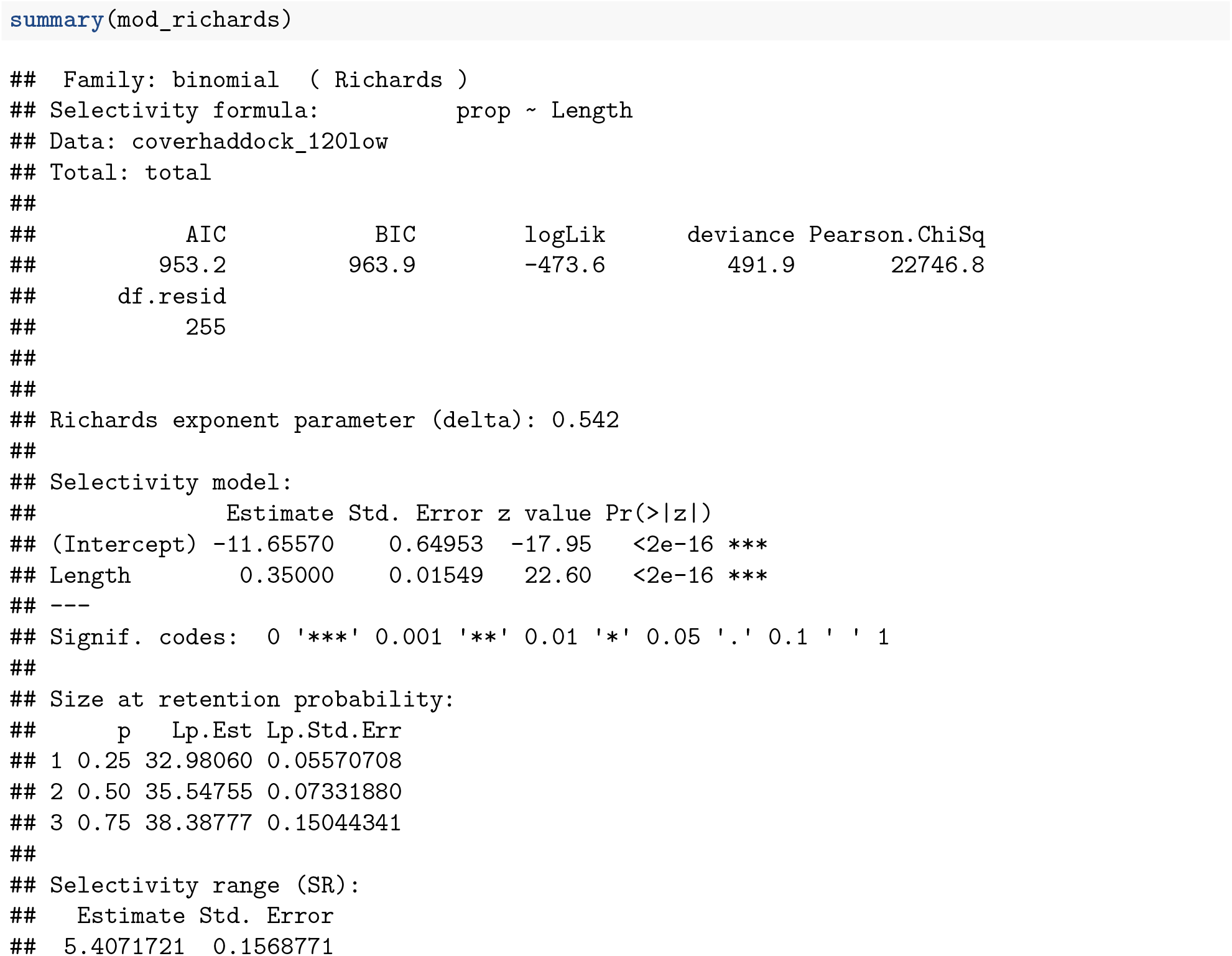

### Predictions

To see how the model fits the data, in addition to residuals as above, it helps to plot observations and predictions together. This could be done to examine any of the models above. We could have compared them using log-likelihoods or information criteria, but we’ll demonstrate how to do that in the next section.

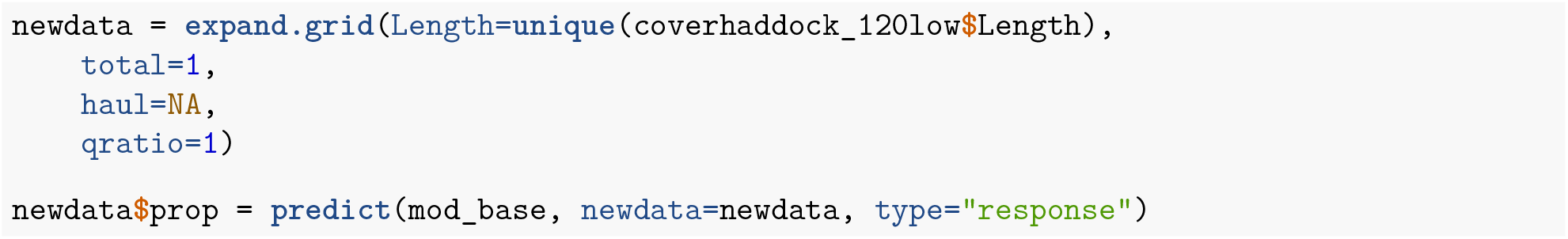

For plotting observations, we need to aggregate the hauls and raise the data by the raising factor. The raised data is only used for plotting, not for statistical analyses.

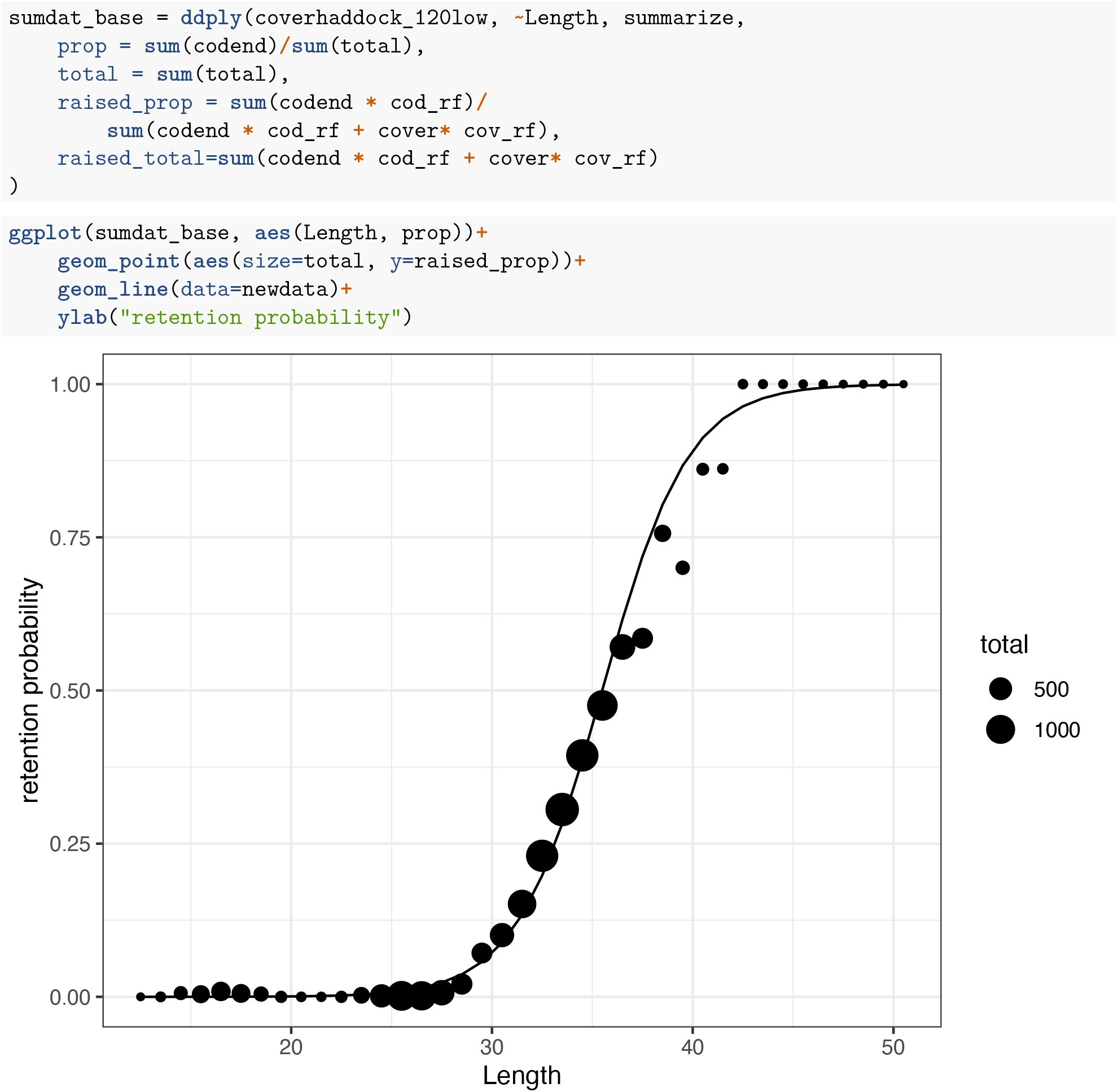

### Confidence intervals by bootstrapping

Following the method of Millar (1994) the bootstrapping function bootSel resamples hauls, then resamples fish within hauls, fits the model to the resampled data, then applies a function `FUN` to each fitted model. In the code below, we define `FUN` to make predictions from each fitted model onto `newdata`. The type of predictions we want in this case are the retention probabilities, i.e. the estimated selection curve, so we specify `type=“selection”`. To read about the `predict` function, type `?predict.selfisher` in the R console.

Mac and Linux bootstrapping in parallel

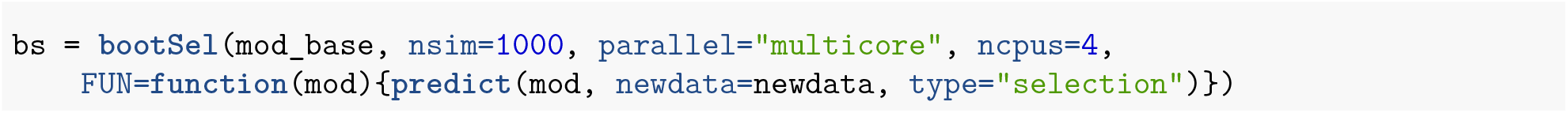

Windows bootstrapping in parallel

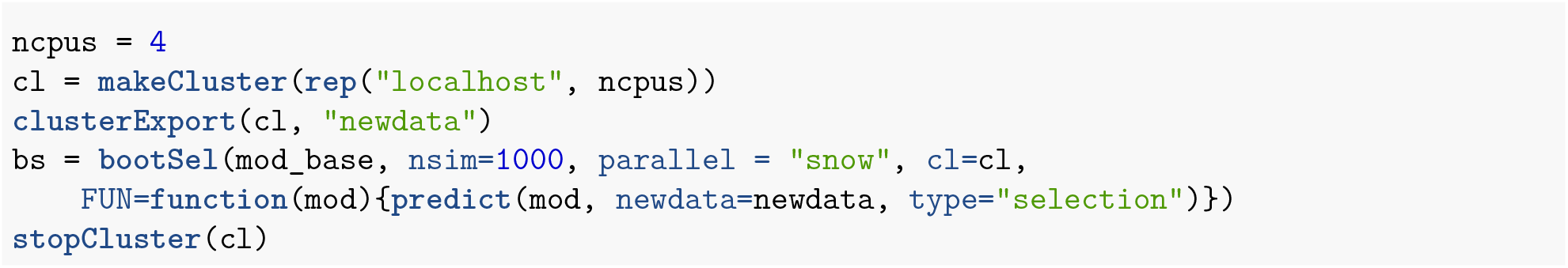

Then we calculate quantiles across bootstraps for each row of newdata.

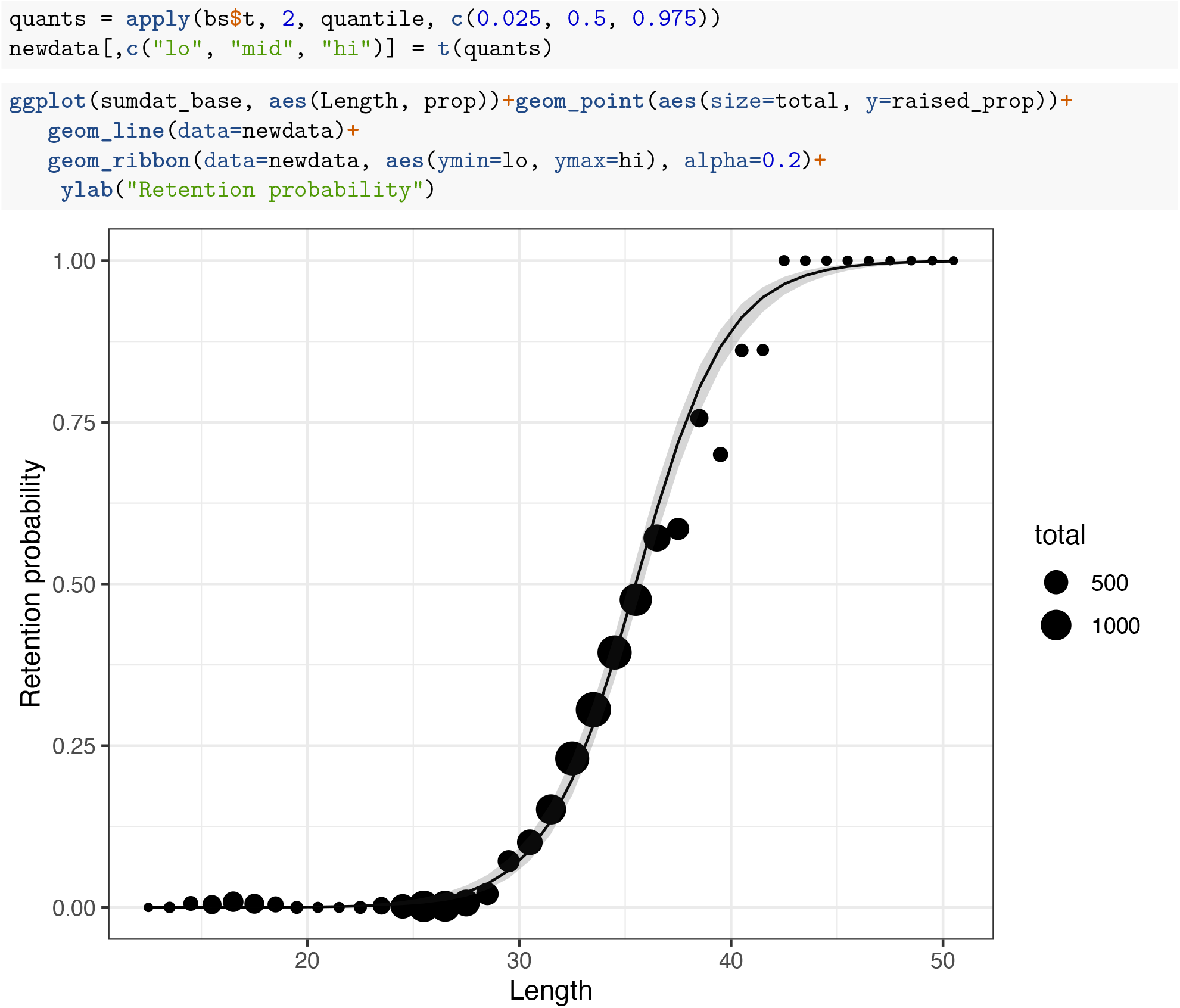

### All four codends in one model

We can analyze the data from all gear types together and test if the gear type affects selectivity by comparing models of varying complexity. If we want to use models directly (i.e. before bootstrapping) for testing the significance of a variable, we need to account for variability among hauls and avoid pseudoreplication (Hurlbert 1984) by including a random effect of haul. The random effect is only needed when models are used directly for hypothesis testing e.g. here, to test our hypothesis that the gear types differ in their selectivity.

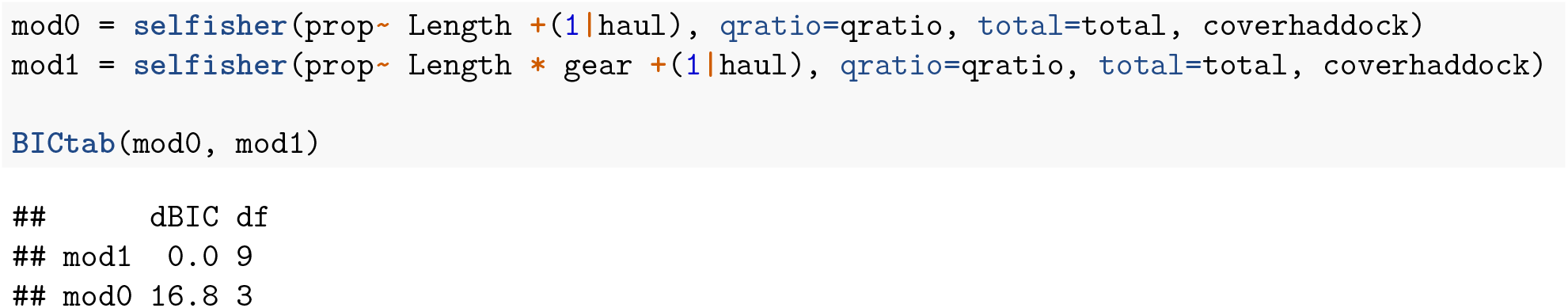

We use the Bayesian information criterion (BIC) to choose the best fit and this table tells us that mod1 is more parsimonious, which means that the 4 codends are somehow different. To explore which aspect of the gears made them different, we examine models with mesh size, twine bending stiffness and catch size. Further, we assume that Baranov’s Principle Of Geometric Similarity applies, i.e. selectivity is a function of fish length scaled by mesh size. It could also be reasonable to fit models with a main effect of mesh size and length in addition to an interaction term, but we do not attempt to fit all possible models here and leave it to users to decide what applies to their particular study.

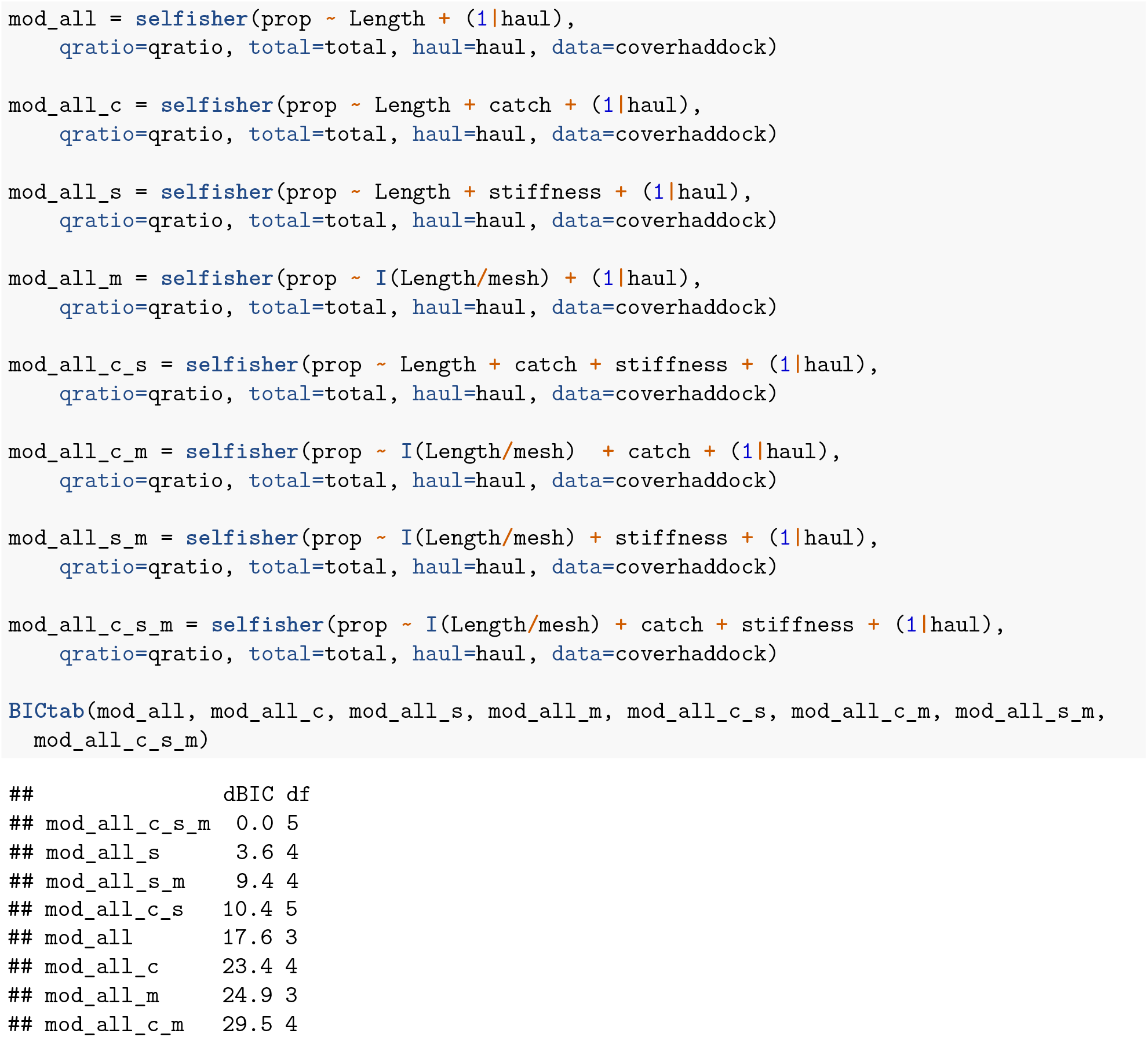

Again we use BIC to choose the best fit and this table tells us that the model that contains mesh size, twine bending stiffness and catch size is the most parsimonious fit to the data.

### Bootstrapping

We drop the random effect of haul from the most parsimonious model before bootstrapping for two reasons (1) the bootstrapping method resamples among and within hauls and thereby accounts for variation among hauls, and (2) random effects make model fitting much slower which can be burdensome when refitting the model 1000 times or more.

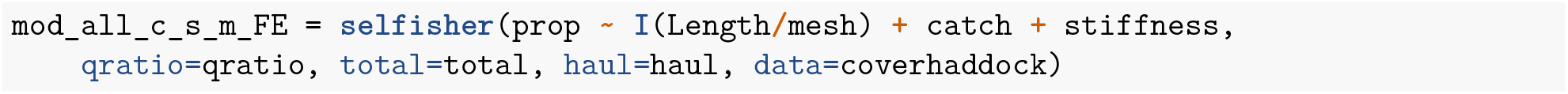

Then we create a new data set to use for predictions. It must include all variables that appear in the model. Even though haul is not used in the mathematics behind the predictions, it must be included in the new data for technical reasons. We include gear just because it makes it easy to organize and plot the data further down. For each gear, we will make predictions over the range of catch weights observed for that gear.

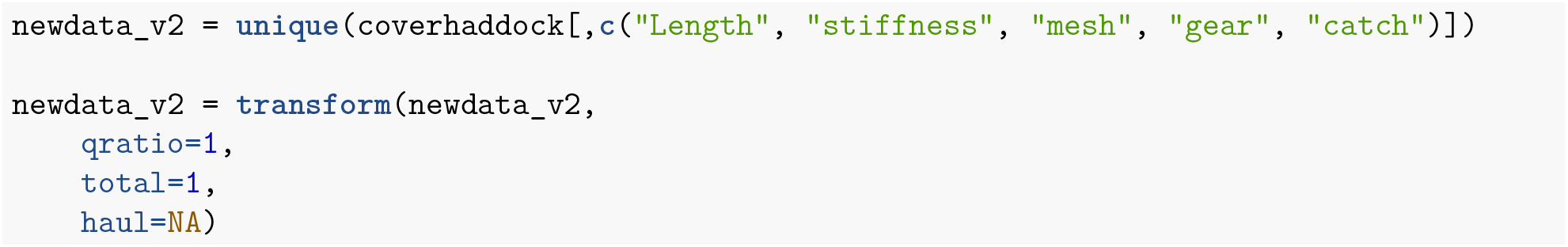

Windows bootstrapping code

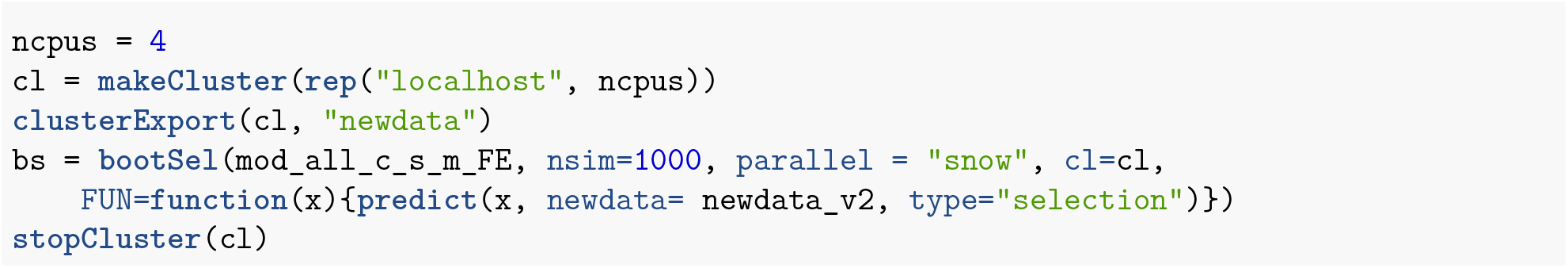

Mac and linux bootstrapping code

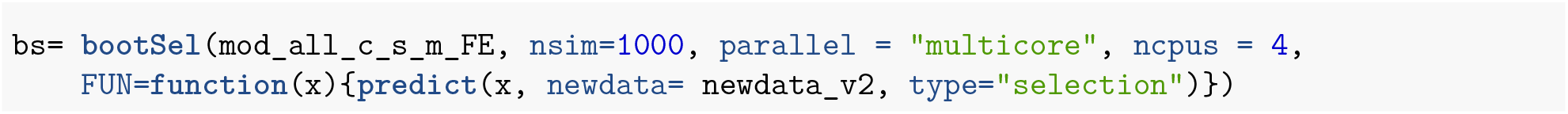

Organize bootstrap predictions with the predictor variables

Then we organize the bootstrap results and join them with the newdata used for predictions.

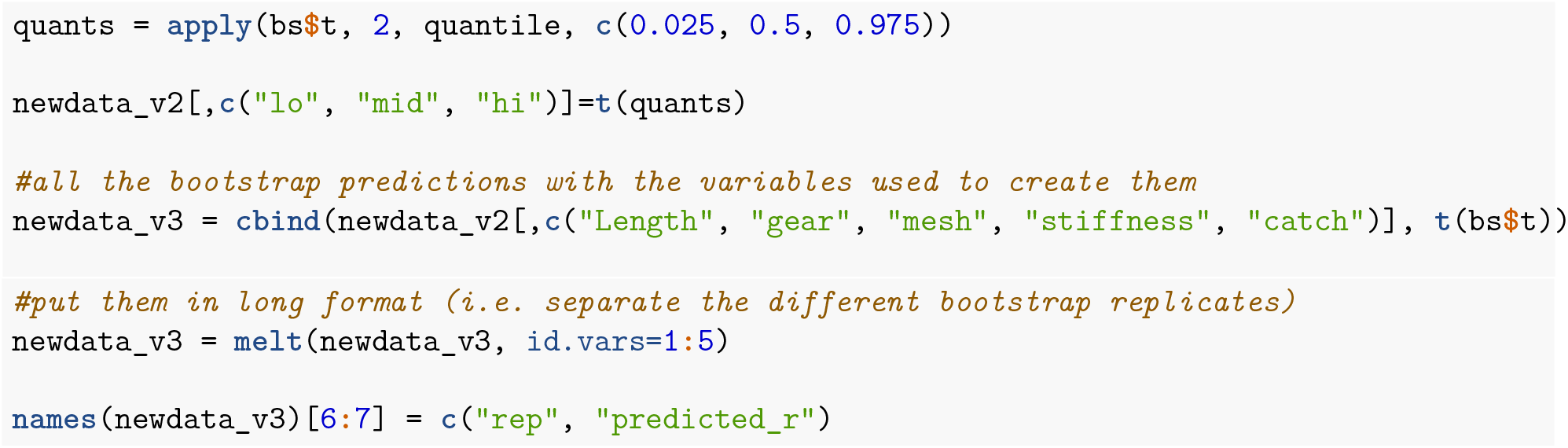

### Calculate *I_50_* and *SR* from bootstraps

Here we define a function to evaluate the length at which a given proportion of fish (p) are retained using interploation.We use this function to find the *l_50_* and *SR* for each bootstrap while varying over mesh size, twine bending stiffness and catch size and subsequently find the mean *I*_50_, mean *SR* and their 95% confidence limits. We plot the results against catch size, which is how they were presented in the original analysis of O’Neill et al (2016).

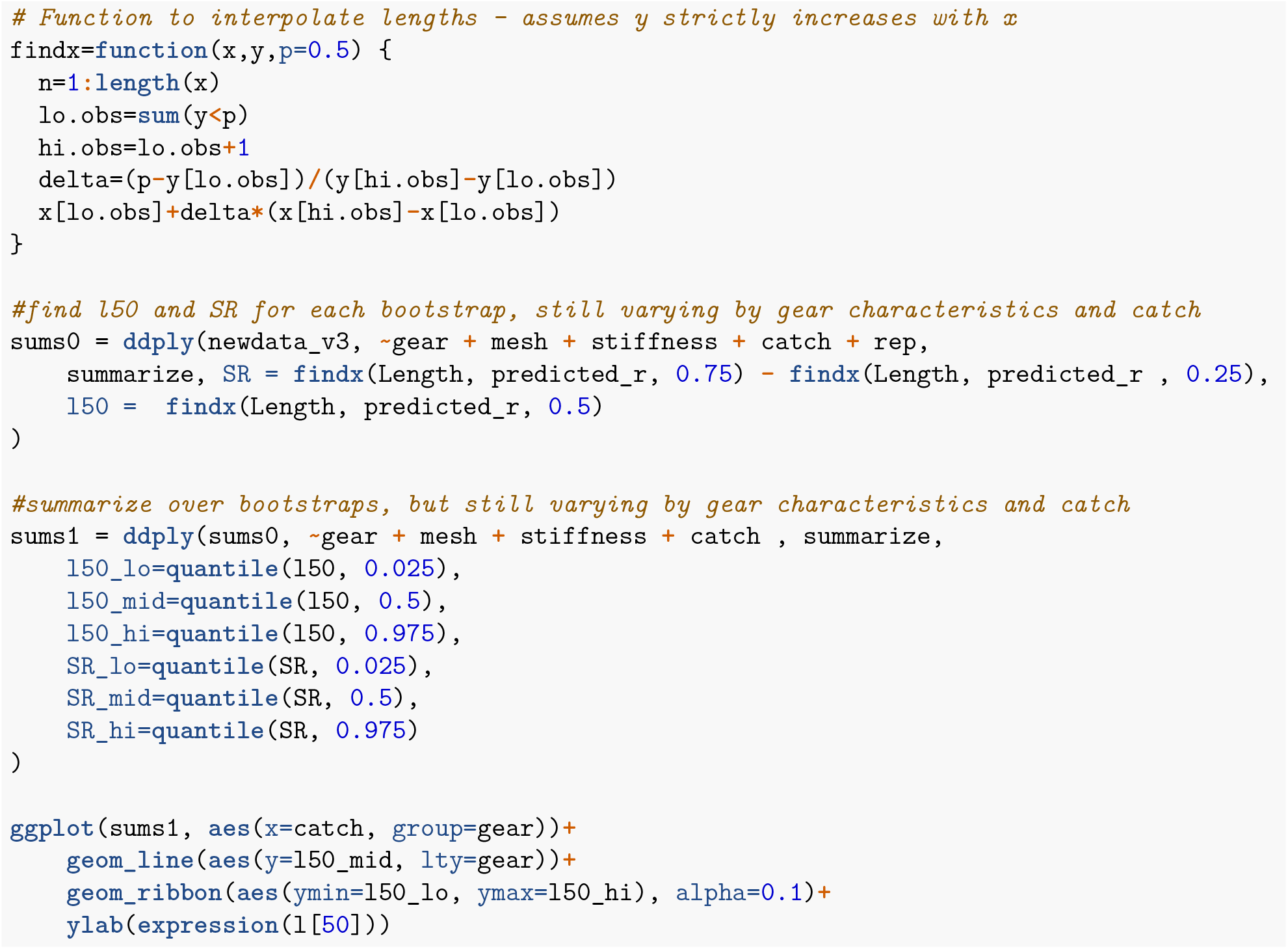
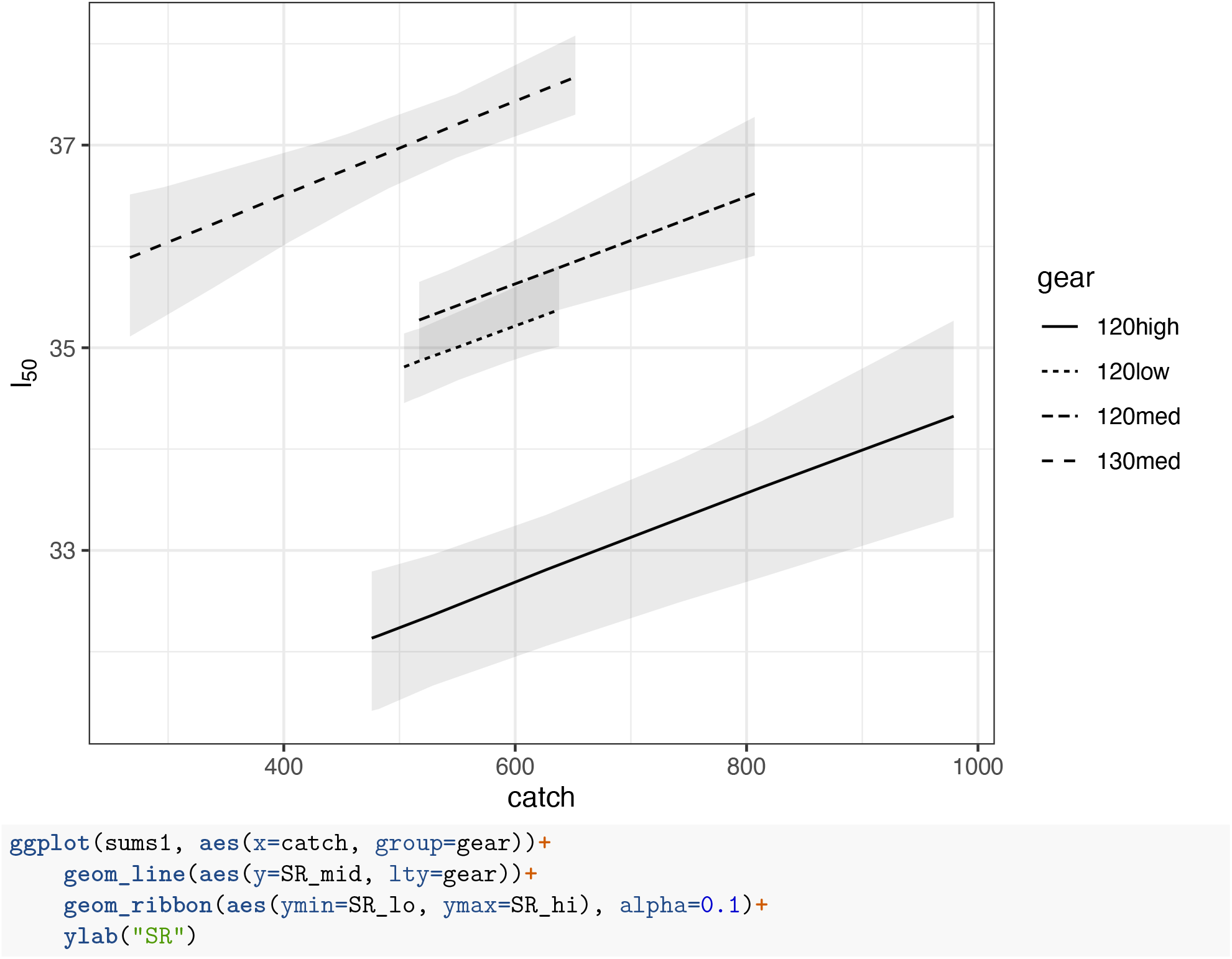
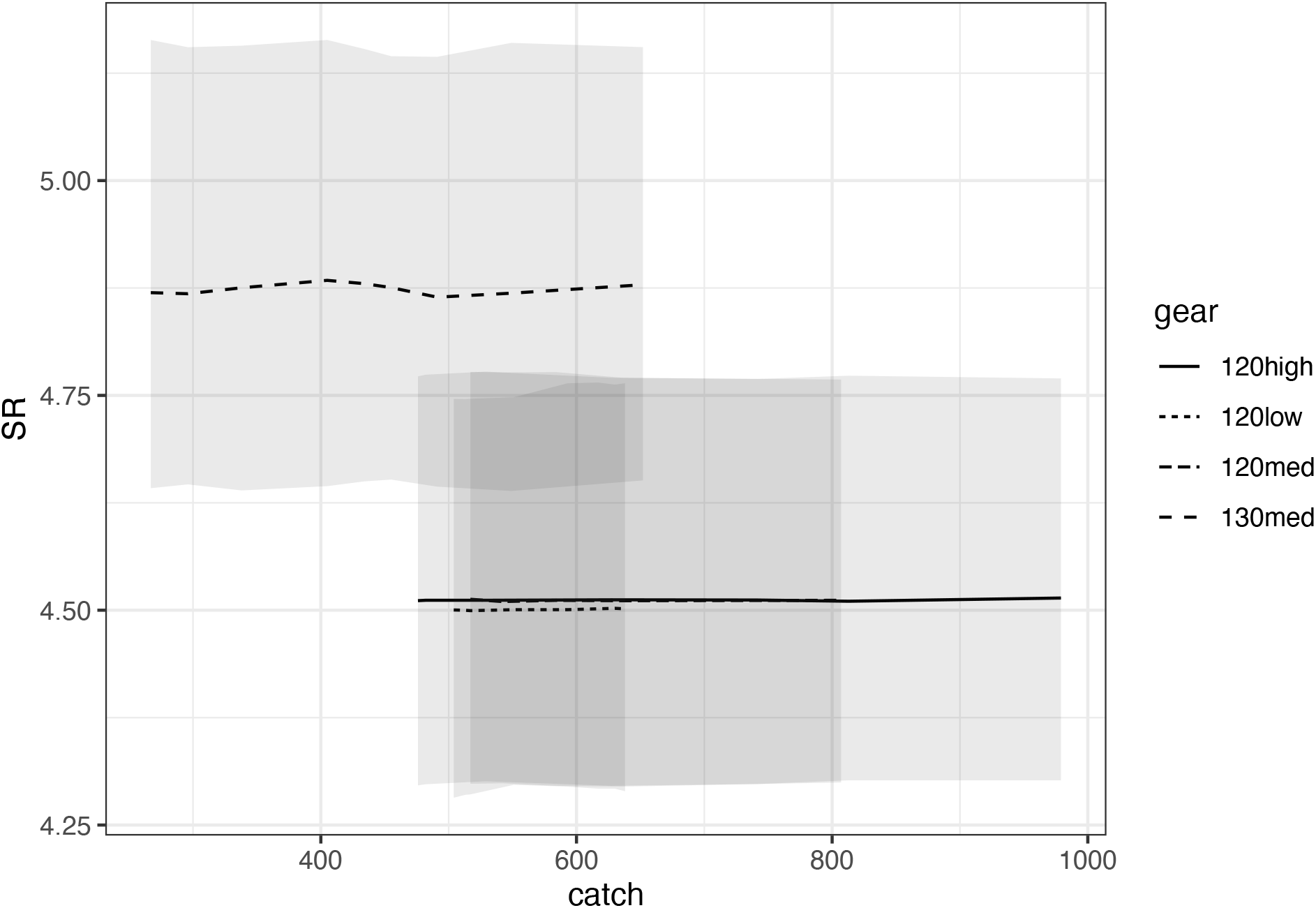

The plots presented here are very comparable to Fig 1 from the original manuscript, which finds a similar dependence of haddock *l_50_* on mesh size, twine bending stiffness and total codend catch weight but finds that *SR* is a constant (O’Neill et al., 2016). We should not, however, expect the results to be identical as there are many differences between the analyses. In the original, it was assumed that the slope and intercept of the logistic link functions vary randomly from haul to haul and that *I*_50_ and log *SR* mean selection curve were linearly dependent on the explanatory variables. Whereas, here we have assumed geometric similarity and that overall retention is related to the explanatory variables.

Below we plot retention curves for each gear aggregated over catches.

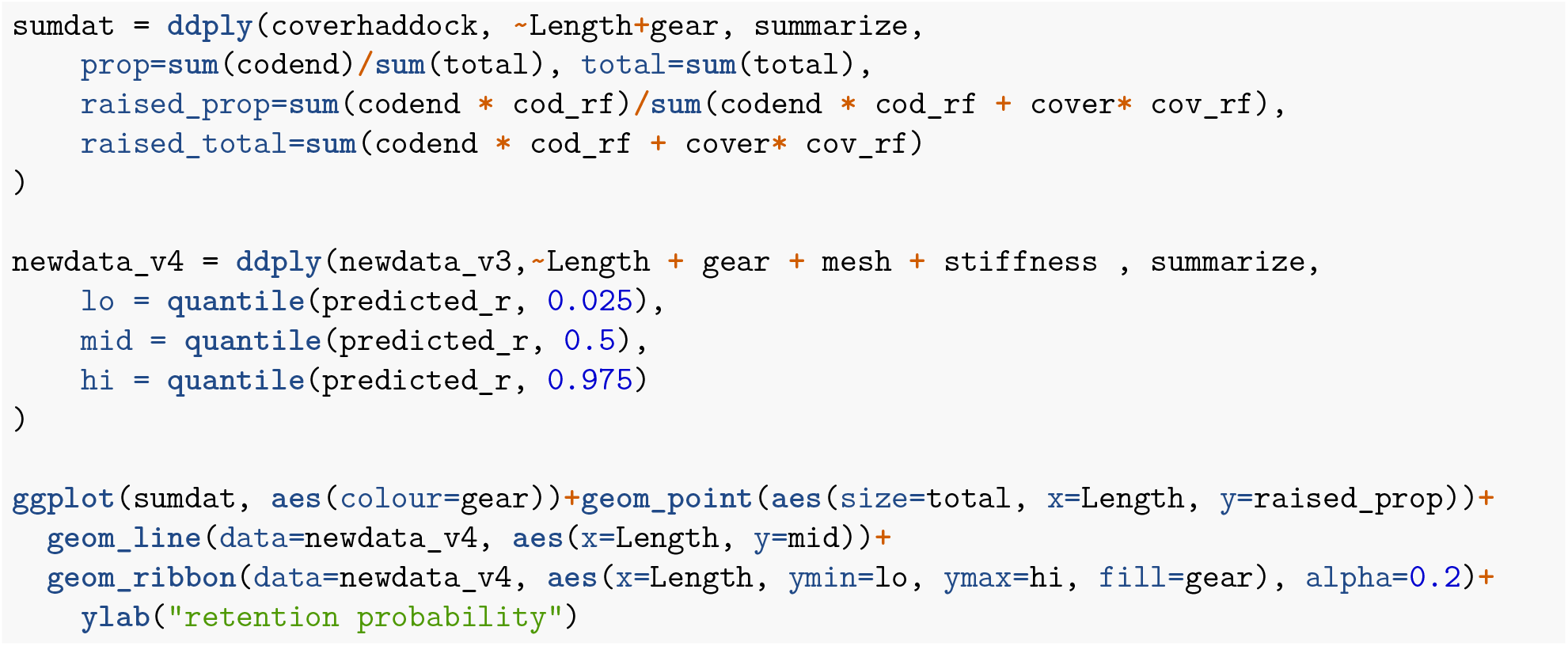
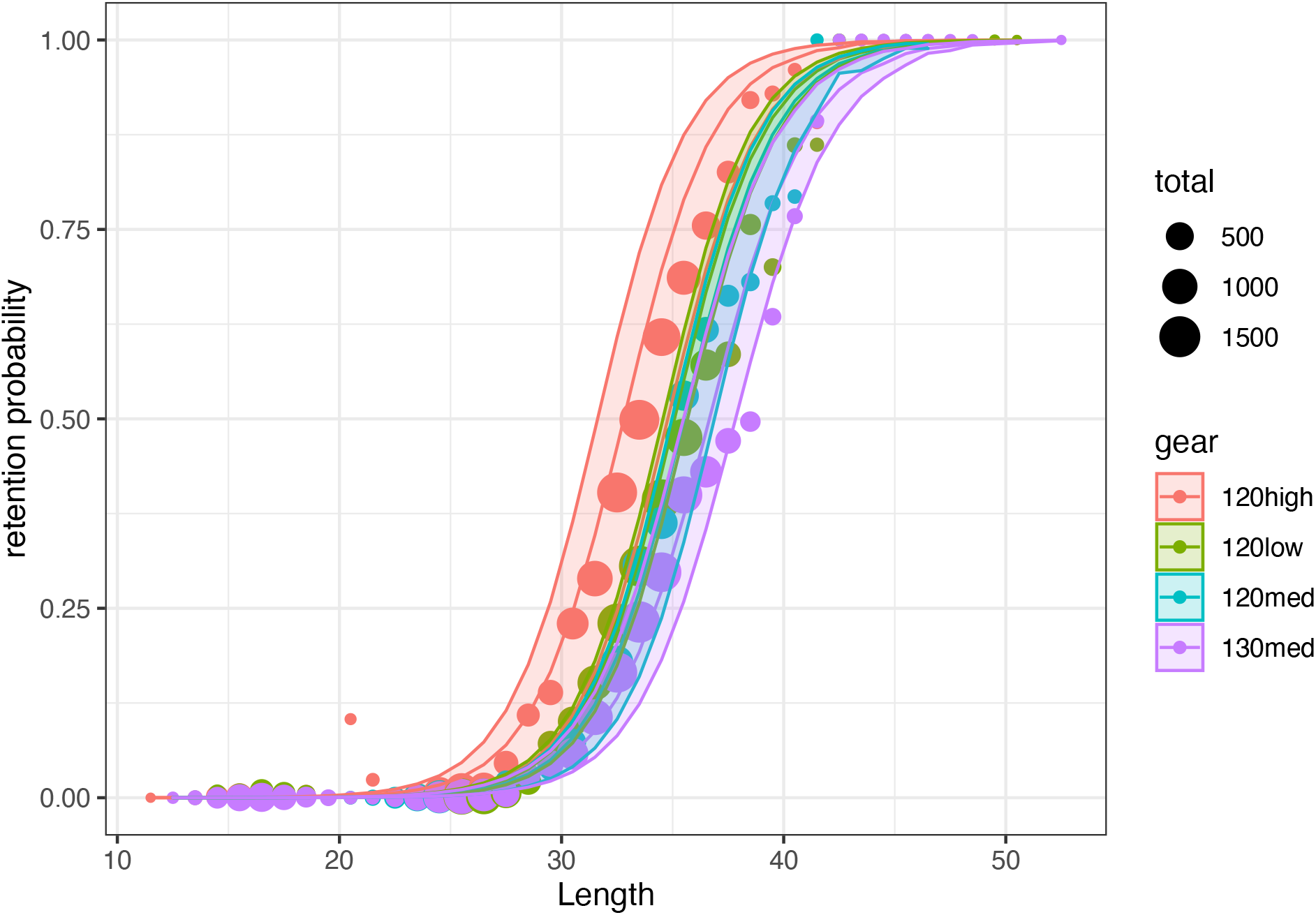

## Appendix 2 Paired gear analysis of codend selectivity dependent on mesh size

### Case study background

This example deals with brown shrimp selectivity data from Santos et al. (2018). The brown shrimp beam-trawl fishery is one of the most important fisheries in the Southern North Sea. Despite its relevance, this is also one of the least regulated fisheries in European waters. Concerns regarding the size selection of the fishery motivated the German research project *CRANNET* (2013-2015), which assessed brown shrimp size selection of commercially used and alternative codend designs. Codend selectivity data was collected during experimental fishing trials using the paired gear method (Millar and Walsh, 1992; Wileman, 1996). The experimental method consisted of fishing with two identical beam trawls, simultaneously and in parallel on the same shrimp population. One of the trawls mounted a small-mesh (11 mm) control codend with very limited selectivity (assumed to be non-selective) on the range of shrimp lengths available for the trawl. The second trawl mounted, one at a time, a total of 33 different test codend designs varying in mesh size and mesh type were tested.

This case study draws on a subset of the *CRANNET* data to demonstrate the use and functionality of `selfisher` in selectivity analysis based on paired gear data. It uses a subset of the catch data, the 87 hauls during which 13 diamond-mesh codends varying in mesh size (ranging from 19.1 mm to 36.3 mm) were tested. Additional information relative to fishing conditions and catch characteristics were recorded at haul level.

### Preliminaries

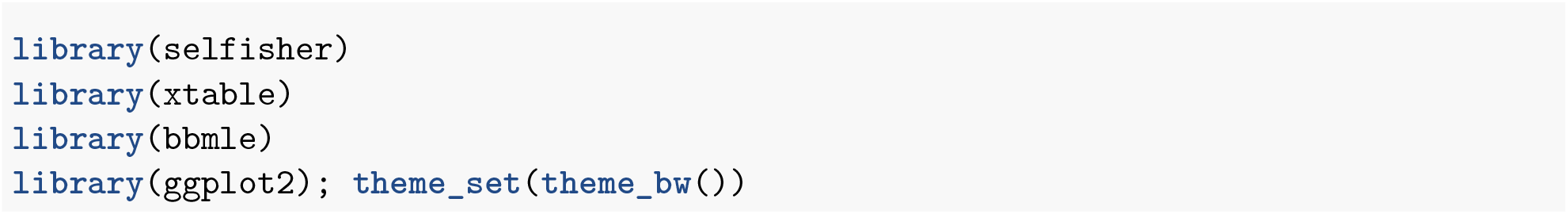

### Data structure

We can read in the data that is distributed with the package and examine it.

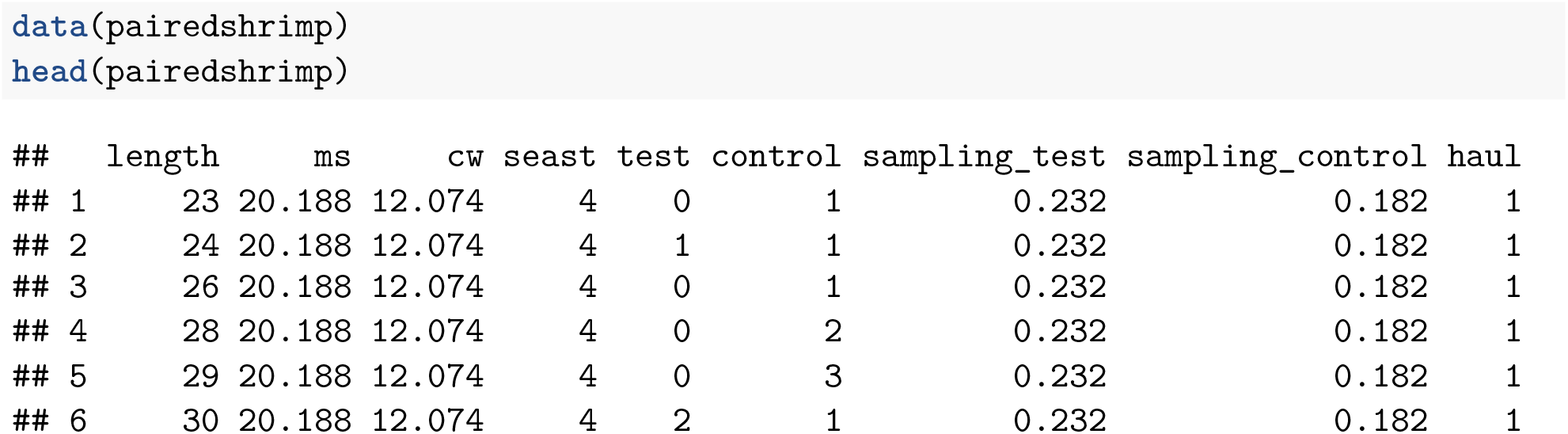

Here we can see the column names of the data. There is a column for haul because multiple hauls *h, i* = 1,2,…, 87 are contained in the same data object. Therefore the information presented in the remaining columns was collected at haul level. The column `length` contains observed (total) length classes *l* of brown shrimp (mm), ms contains the measured mesh size (mm) of the test codend, cw is the total catch weight (kg) collected in the test codend, `seast` is the state of the sea recorded during towing (Beaufort scale). Columns `test` and `control` contain the numbers of shrimps of length *l* sampled from each of the paired codends, while `sampling_test` and `sampling_control` are the associated sampling fractions.

### Data transformation

In `selfisher`, it is necessary to model the binomial response as a proportion and total because unlike many other methods for binomial GLMs, the underlying code does not accept a two column response variable. Therefore, we transform the data to create these new columns. We also create a column for `qratio` (i.e. *qh* below) which is the ratio of the sampling fraction of the test over the control gear. If the data contained raising factors instead, then `qratio` would be the raising factor of the control gear over the test gear.

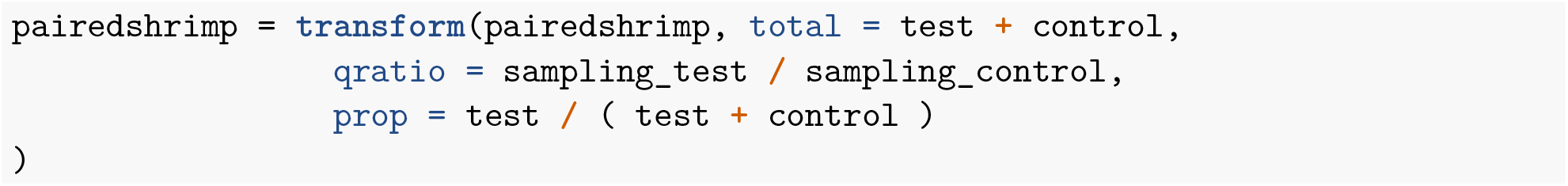

### Selectivity analysis

The selectivity analysis based on paired gear data is usually done with the model introduced by Millar and Walsh (1992):

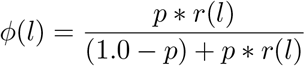

In the equation above, *φ(l)* expresses the probability that a shrimp of length *l* in the total catch of the paired gear was caught by the test gear. This probability is related to a sequence of two fishing events. The first event is controlled by the probability that a shrimp entering the paired gear did it through the test trawl (relative fishing power), expressed as the length-independent split parameter *p*. Conditioned on the probability of entering the test trawl, the second event is the length-dependent retention probability of the test codend *r(l)*. Retention probability is usually (but not exclusively) modelled with the logistic function, and summarized by two selectivity parameters *L*_50_ (length of 50% retention probability) and *SR* (difference in the lengths of 75% and 25% retention). Further details of this model and extensions involving alternatives to the logistic function can be found in Millar and Walsh (1992) and Wileman (1996).

In Santos *et al.* (2018), the effect of codend design as well as other variables describing catch and operational characteristics of the hauls were assessed using the so-called Fryer method (Fryer, 1991). The Fryer method is carried out in two steps. In the first step the parameters *l_50_, SR, p* and associated Hessian-based covariance-variance matrix of individual hauls are estimated. The estimates become the data used in the second step, where the effect of the measured explanatory variables (fixed effects) on codend selectivty is estimatated. Such estimations are obtained using the EM-algorithm, which allows quantifying the strength of the fixed effects in the presence of between-haul variation.

The Fryer method was developed at a time before suitable generalized mixed modeling software such as `selfisher` was available. The Fryer method suffers from small sample bias in the fits to individual haul data. In contrast, `selfisher` enables quantifying and testing the effect of explanatory variables in a single step and directly on the measured catch data.

The methods in `selfisher` generalize the original selectivity model introduced by Millar and Walsh (1992), by allowing multiple fixed and random effects to simultaneously model relative fishing efficiency and the selectivity of the test codend:

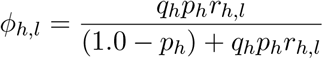

In the equation above, 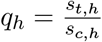 is the ratio of the fraction of brown shrimp sampled in the test gear

to the fraction sampled in the control gear, *p_h_* is the relative fishing power of the test gear in haul *h* (split parameter), where 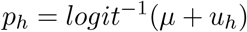 and *u_h_* is a random effect 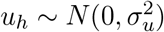 accounting for haul-specific random variation from the mean value on the logit scale *μ*. For simplicity in this example, we assume a logit link in the retention model and therefore the retention probability model for haul *h* and length *l* is 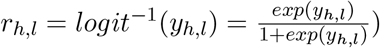. This expresses the haul-specific probability for a fish entering in the test gear to be retained where *yh,ι* is the expectation on the link scale combining fixed- and random-effects potentially influencing retention probability of the test codend. Four models varying in the structure of *y_h,l_* are initially considered:

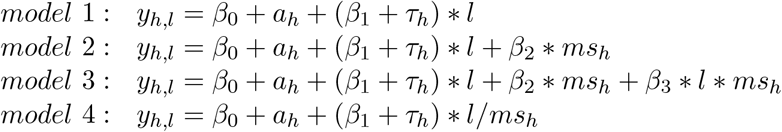

All models listed above include an intercept *β_0_* which expresses the baseline shrimp retention log-odds when all additional covariates (including shrimp length) are set to 0. The coefficient *a_h_* is a random effect 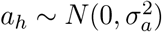 accounting for haul-specific random variation of the intercept. Model 1 assumes that codend retention can be exclusively described by shrimp length. In models 1, 2, and 3, *β_1_* is the slope quantifying the rate of increment in retention probability (on the link scale) due to increments in the length of shrimp. The slope of size selection curves can vary significantly and randomly between hauls, even while keeping the design characteristics of the codend constant (Fryer, 1991). The coefficient 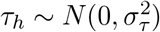 is introduced to account for haul-specific random variation of the baseline *β*_1_ value. Model 2 extends model 1 by adding mesh size (*ms*) as a fixed effect that varies by haul: *ms_h_*. Model 2 assumes that retention probability is a result of separate effects of shrimp length and mesh size. With the addition of an interaction term of mesh size and shrimp length (*γ*_3_), in model 3 it is assumed that mesh size influences both the position and slope of the size selection curve.

The tested diamond-mesh codends were made of the same netting material, same length, and the number of meshes in circumference were reduced proportionally to increments in mesh size. Therefore, we assume that the meshes of the tested codends present the same geometry during fishing. Based on the principle of geometrical similarity (Baranov, 1948; Millar and Holst, 1997; Revill and Holst, 2004) it would be a reasonable *a priori* assumption that the selection curves will vary systematically with mesh size. Model 4 is built upon the hypothesis that variation in size selection can be explained by the principle of geometrical similarity, implying that the size selection varies proportionally to mesh size (i.e. doubling the mesh size implies doubling the values of *I*_50_ and *SR* of the selection curve)(Millar and Holst, 1997).

### Model fitting with selfisher

In this section, the models described above are fit using a function named `selfisher` within the `selfisher` package. Formulas in these models follow the convention of the `lme4` and `glmmTMB` packages. The function takes a formula for the retention model `rformula` as the first argument. This is a two sided model with the proportion on the left side and fixed and random effects on the right side. For example, in model 1 (m1) below, `prop ~ length + (length | haul)` says that the proportion (`prop`) of fish retained in the test gear depends on length and that the intercept and slope vary randomly by haul. It also takes an argument for the relative fishing power model (`pformula`) which is a one-sided formula, e.g. `pformula =~(1 | haul)`; this says that relative fishing power should be estimated and can vary randomly by haul. If instead, we wanted to fix the split at 0.5, then we could specify `pformula =~0`. To tell the funciton that this is a paired gear model with one non-selective gear, we use the argument `psplit = TRUE`. The function also takes as arguments the names of the columns for the total and qratio within the data frame specified by the argument `data`.

One optional argument appears in this example: the `start` argument. It tells the function what starting values to use in maximum likelihood estimation. See `?selfisher` for the full flexibility of how to specify starting values, but here we only give starting values for the retention model. To get good starting values for the retention model’s intercept (*β_0_* above) and coefficient on length (*β*_1_ in models 1, 2, and 3 above), we use the inits function which takes guesses for *I*_50_ and *SR* as its arguments (30 and 8 respectively here). The coefficient *β*_1_ has a different meaning in model 4, but the starting values work as supported by plots below, so it’s not a problem. Models fit with TMB (as in `selfisher`) are usually robust to starting values, but due to the complexity of paired gear models, they sometimes get stuck in local minima during maximum likelihood estimation. In this case study, the length of the start vector must equal the number of fixed effects coefficients in the retention model, i.e. the *β*s in the equations above, so in m2 and m3 we combine guesses for *β*_0_ and *β*_1_ with zeros for the other coefficients

As described above, model 4 (m4) assumes geometric similarity. To include *length/ms* in a model in R, it is necessary to tell the formula interface to use the term “as is” by putting an *I*() around it.

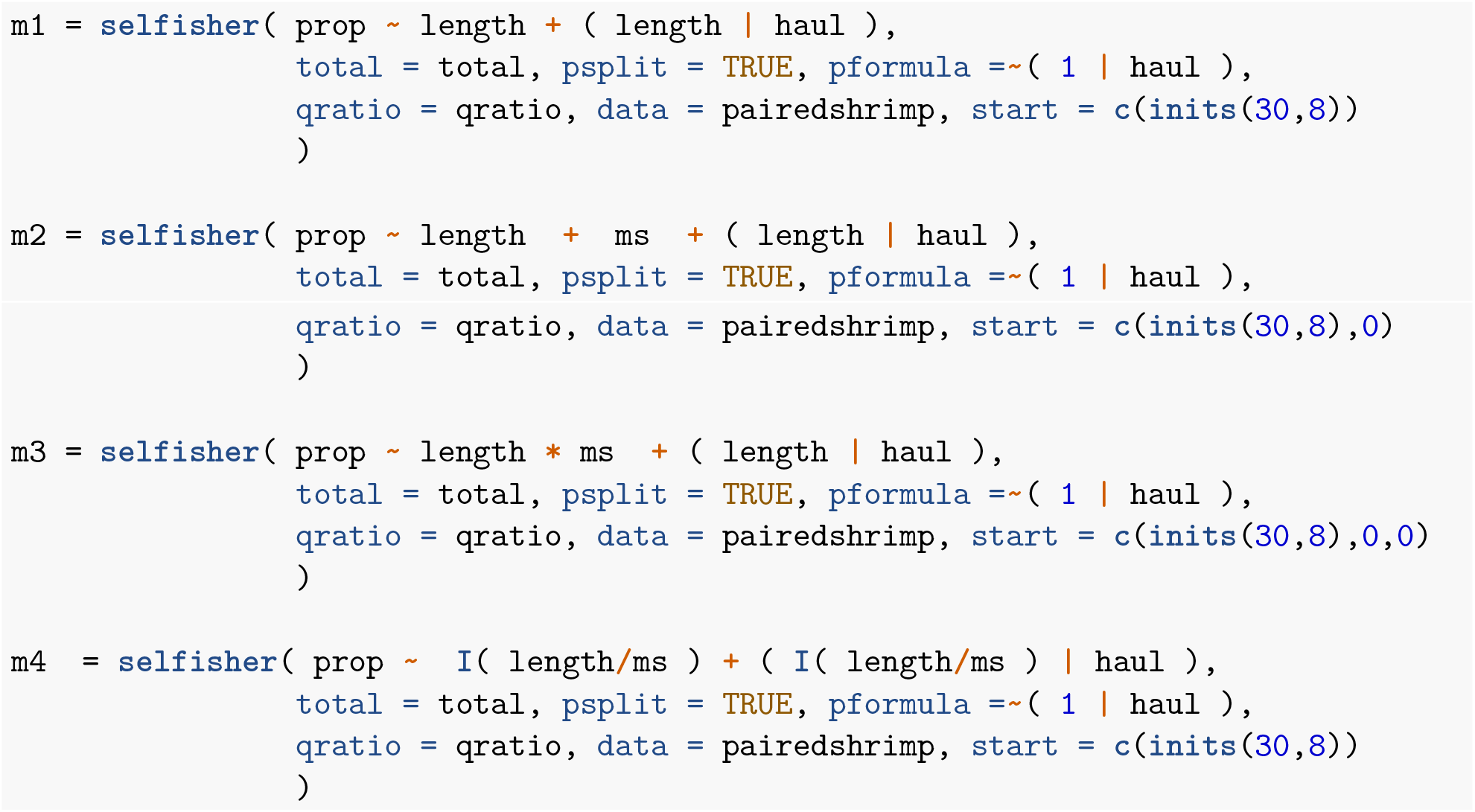

### Model selection

Retention is a mechanical process that can be largely explained by the relationship between the mesh characteristics and the morphology of the species being selected. Of the above models, model 4 is the only one that has a mechanistic justification, namely geometric similarity. In terms of model selection we *a priori* propose model 4, taking the view that strong evidence against model 4 is required to prefer an alternative. There are several thousand observations in the data, and consequently Akaike Information Criterion will tend to choose the most complicated model (Heinze *et al*., 2018). In contrast, the Bayesian Information Criterion, BIC, includes a stronger penalty for the number of parameters than AIC and therefore it tends to select simpler models than AIC. We can use functions from the bbmle package to create either an AIC or BIC table of the models.

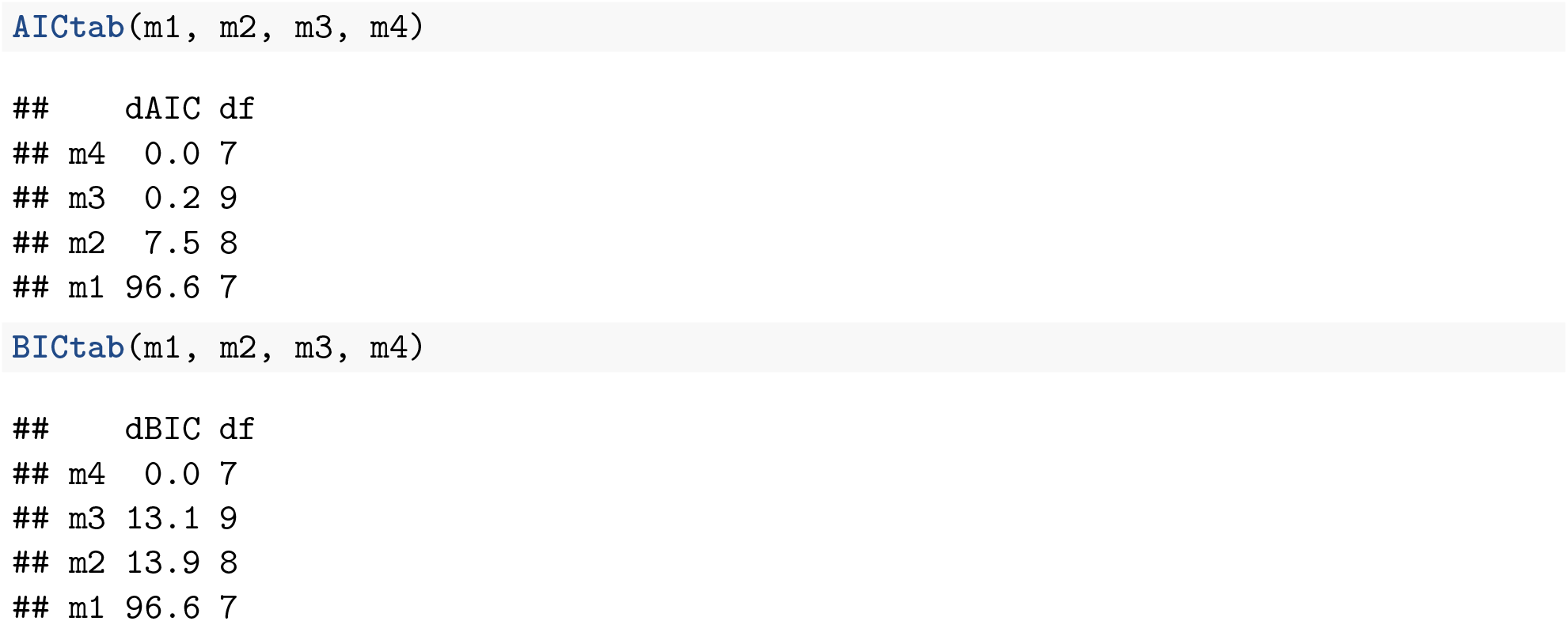

Both AIC and BIC rank model 4 on top, as was the *a priori* expectation.

### Extended models

Model 4 is now extended by adding other covariates in the data frame that could potentially influence size selection, such as cw and seast. The following models are fitted following the same procedure as for models 1 to 4:

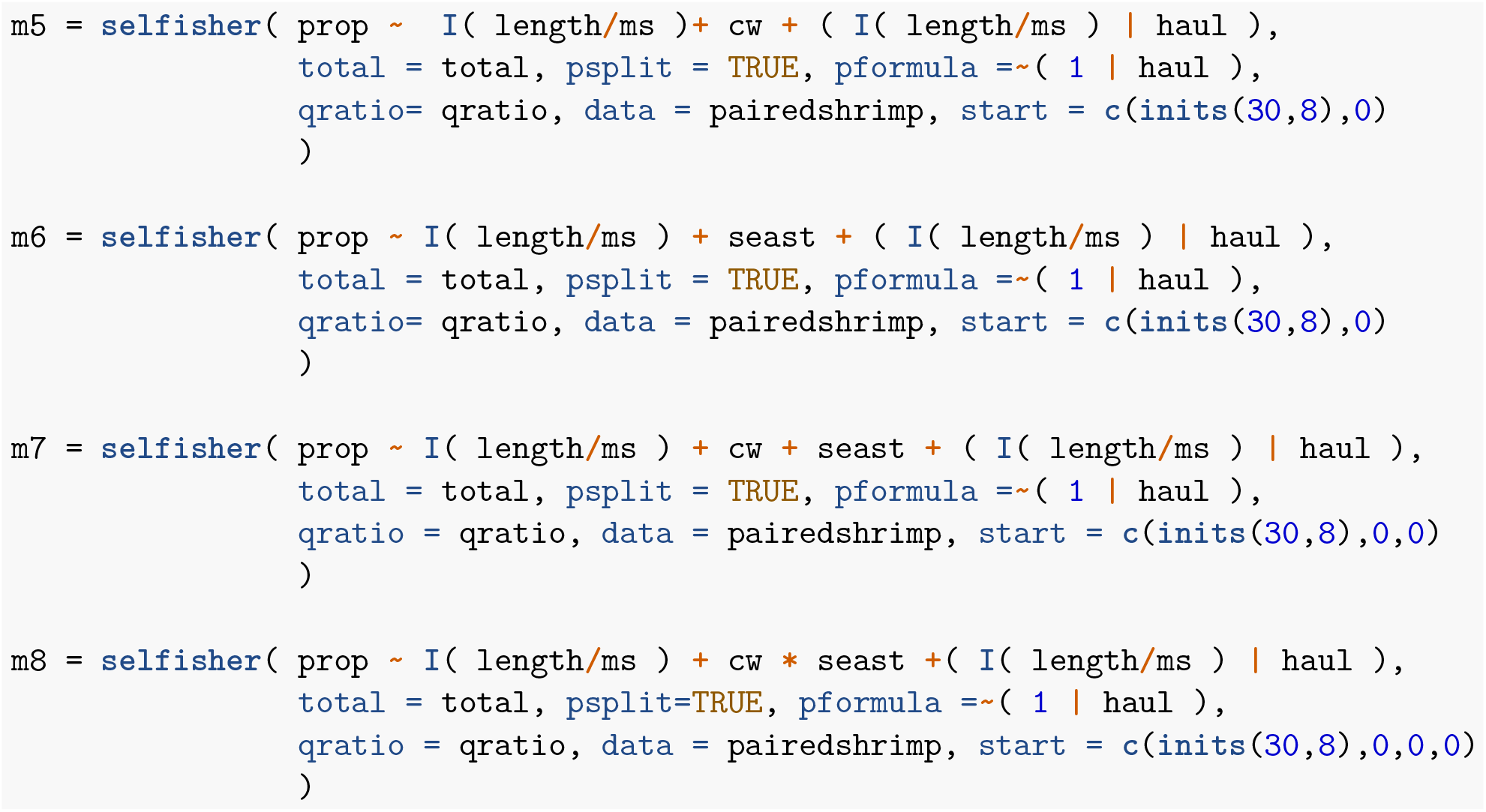

Then we can compare all the models.

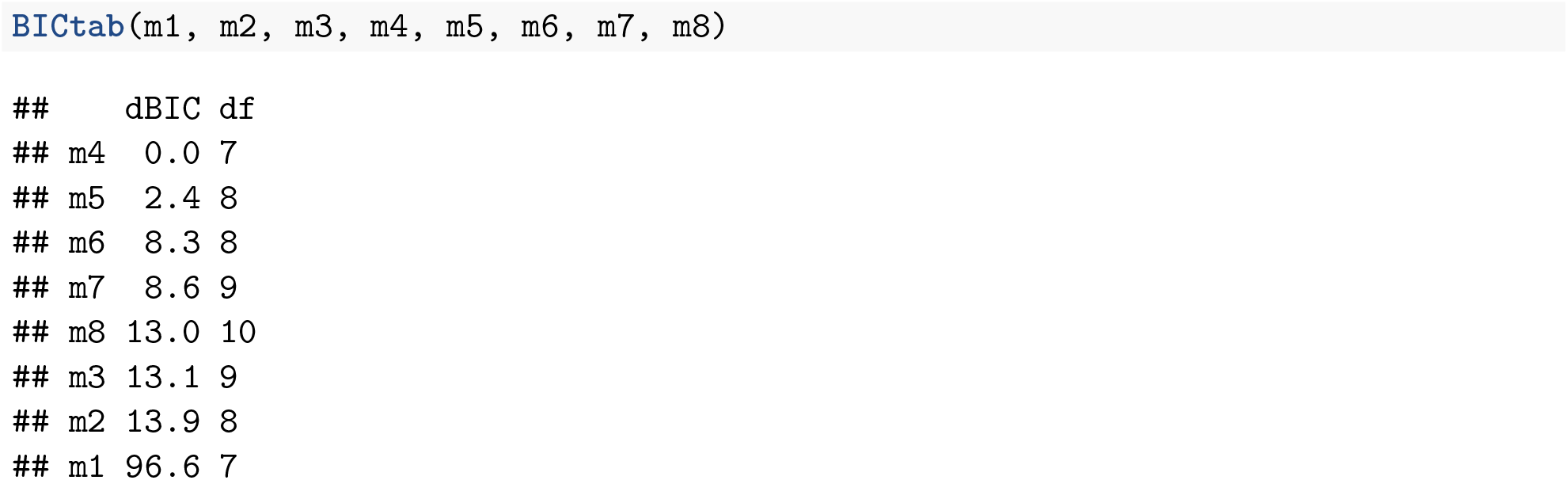

This table shows that none of the individual or combined effects associated with cw and `seast` were strong enough to be included in the most parsimonious model. Therefore model 4 is selected in this case study for further assessments.

### Simple results

Consistent with other model procedures implemented in R, a summary of model results and fit statistics can be obtained via `summary(model.object)`. Before inspecting the size selectivity results provided by model 4, the estimated relative fishing efficiency of the test gear is presented. By default, `selfisher()` summary shows the estimated split parameter *p* on the logit scale, but this might be updated in a new version. An inverse logit transformation is needed to obtain the fishing power *p* ∈ [0,1]:

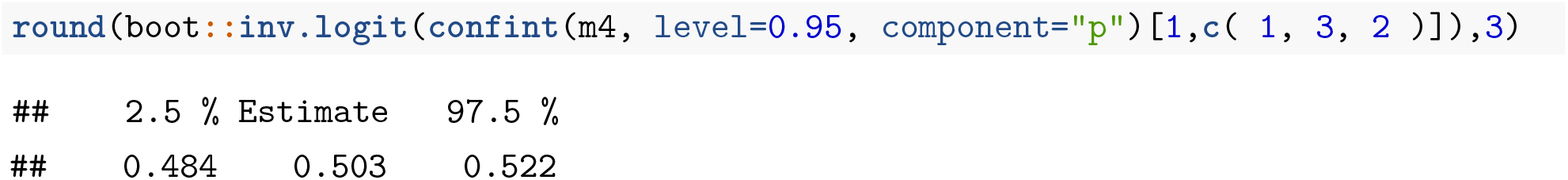

The split parameter estimated by model 4 is *p* = 0.503 with 95% confidence interval (0.484 — 0.522), very close to the value estimated in the original study (*p* = 0.492 (0.472 — 0.512)). This includes the reference value of 0.5, from which we conclude that there is no significant evidence against equal catch efficiency of the test and control gears.

### Goodness-of-fit

The mixed models presented above are fitted at haul level. Therefore it is reasonable to assess the goodness-of-fit of these models on individual haul data. Due to the large dataset used, a random sample of 12 hauls are picked to demonstrate how well the *φ(l)* curves estimated by model 4 fit to the data. We can use the function `predict` to examine the estimated retention curve, ie. the model’s “response” variable. See `?predict.selfisher` for details of how to use this function including the different types of predictions available.

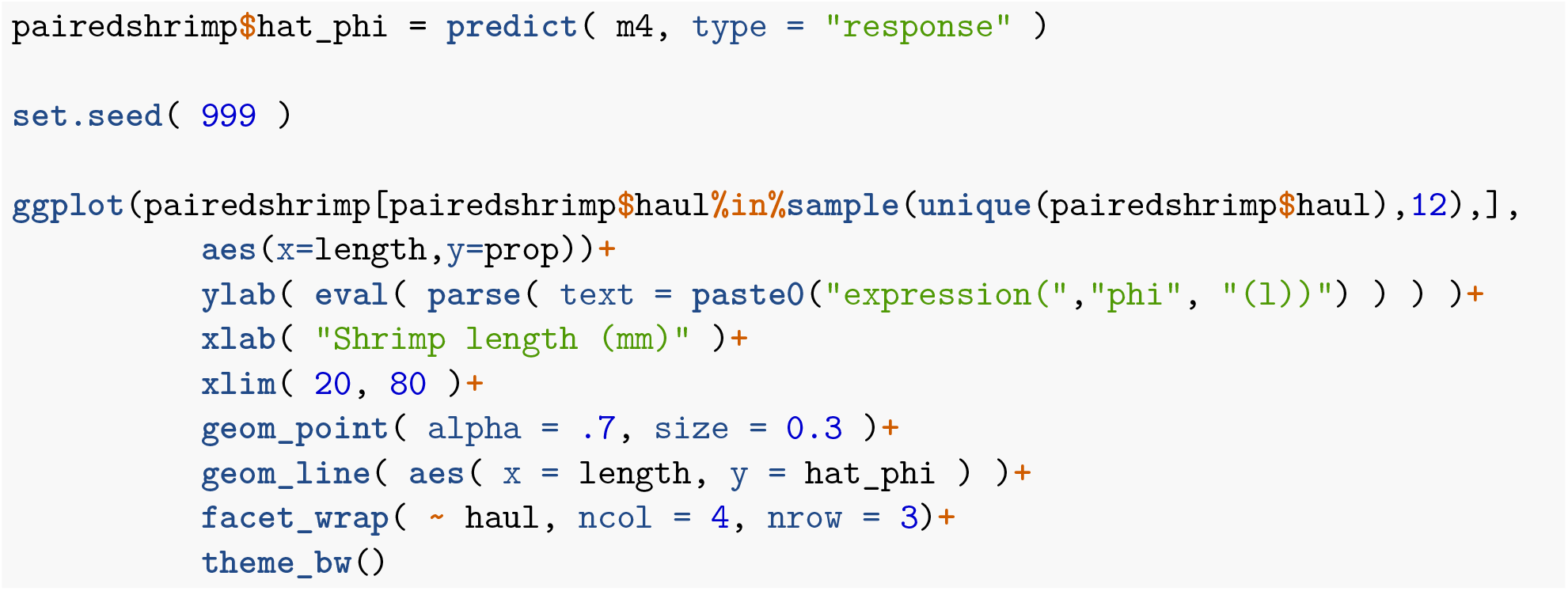
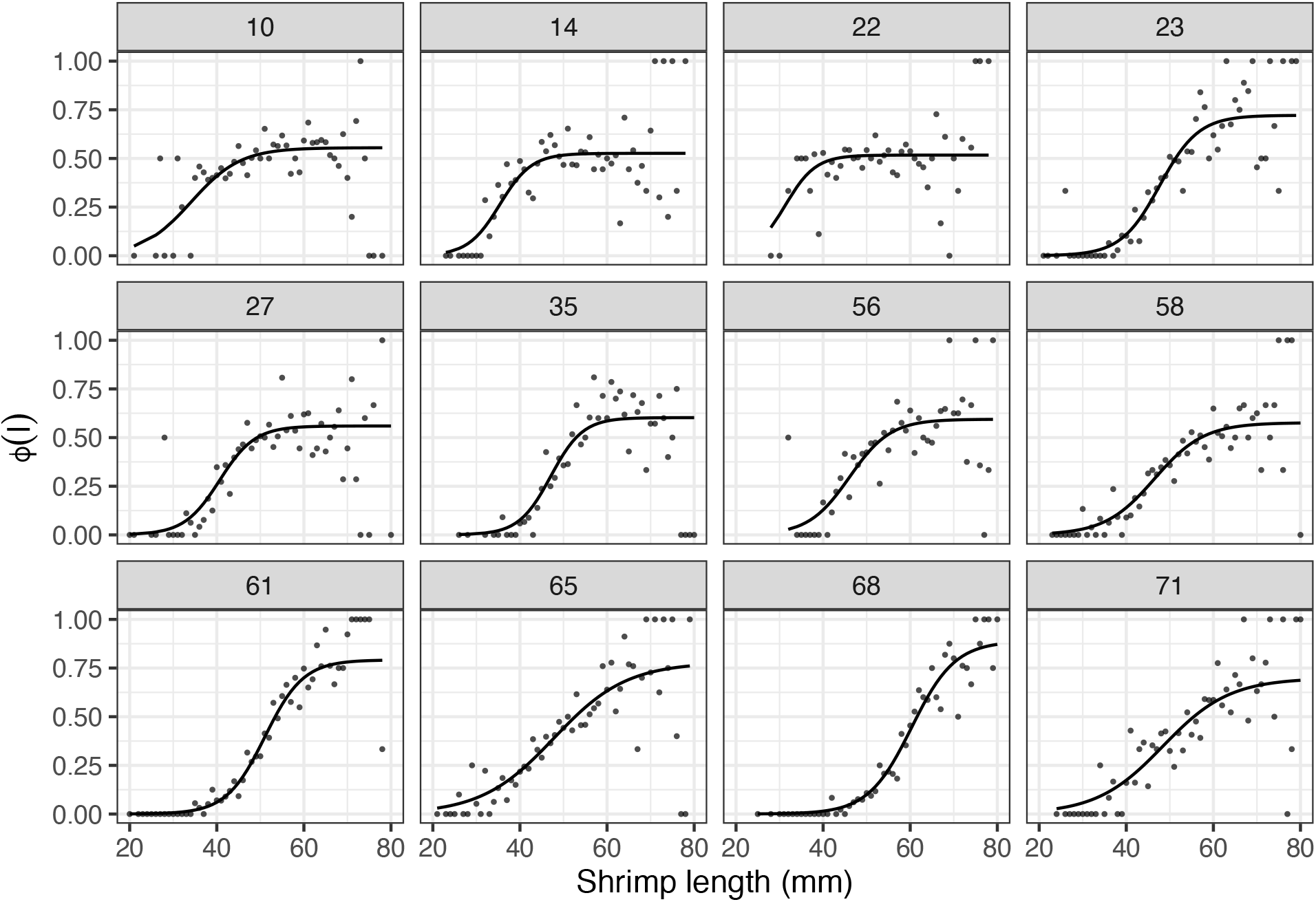

The figure above reveals that the fitted *φ(l)* curves describe well the trends and variability of the data at individual haul level.

### Population-average predictions

Mixed modelling is a formal procedure that takes into account specific details of the data collection enabling sound hypothesis testing on fixed effects and model selection. However, fitting mixed models can be a computational-intensive task. Moreover, the researcher is also typically interested in obtaining average selectivity predictions, as these are relevant to the selectivity applied to the fishery. It is therefore recommended to refit the best candidate model leaving out the random effects. Bootstrapping can then be used to obtain valid standard errors and confidence intervals for estimated quantities such as *I*_50_ and *SR*.

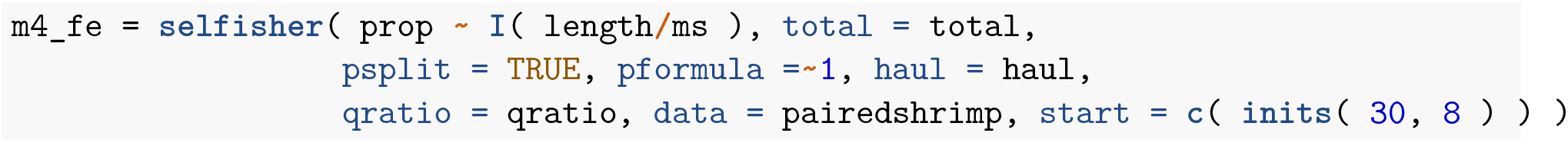

A summary of model coefficients describing codend retention can be obtained using standard procedures in R

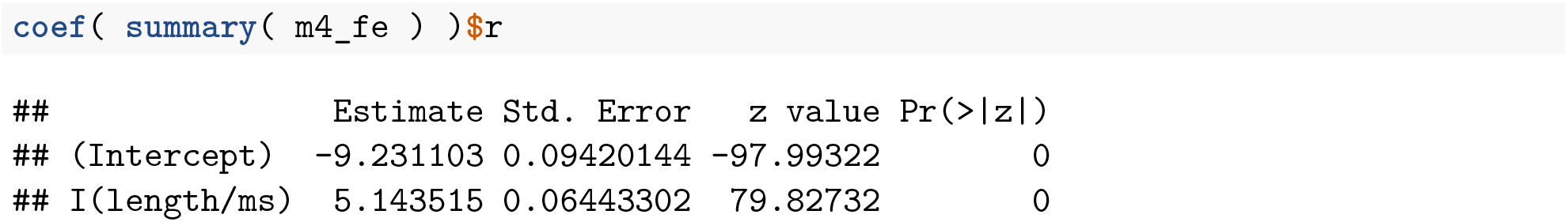

### Bootstrapping to get confidence intervals on population-average *I_50_* and *SR*

Selectivity statistics *I*_50_ and *SR* are often obtained by simple calculations involving model coefficients *β*_0_ and *β* 1, or for simple models in `selfisher`, the function `L50SR(model.object)` can calculate them. However, standard calculations need to be updated when using multiple fixed effects to describe codend retention. Details on how to calculate *I*_50_ and *SR* from the models considered in this case study can be found in Table 1 below.

To obtain a bootstrap distribution of the selectivity parameters estimated by model 4, first we generate a bootstrap distribution of model coefficients using `bootSel()`, as follows. The `bootSel()` function applies the user-defined function `FUN` to each refit model; here we define FUN so that it returns the fixed effect (`fixef`) coefficients of the retention model ($r). This is the code to perform the computations in parallel on Linux or Mac computers, but see the other case studies for how to do it in Windows.

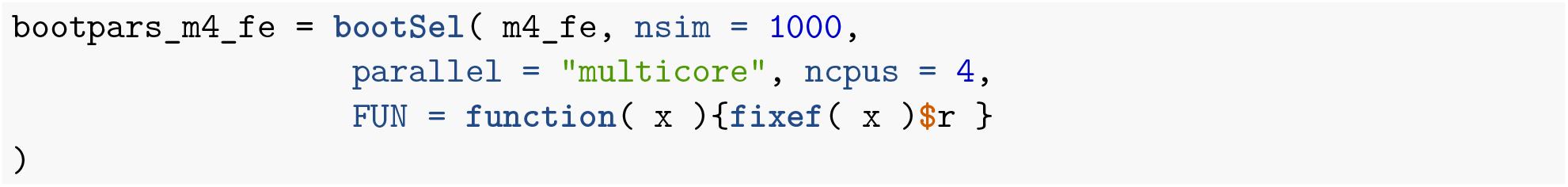

Selectivity statistics *I*_50_ and *SR* are then calculated from each set of coefficients in the bootstrap distribution generated above. Finally, the resulting bootstrap distribution is used to obtain 95% percentile confidence intervals of *I*_50_ and *SR*.

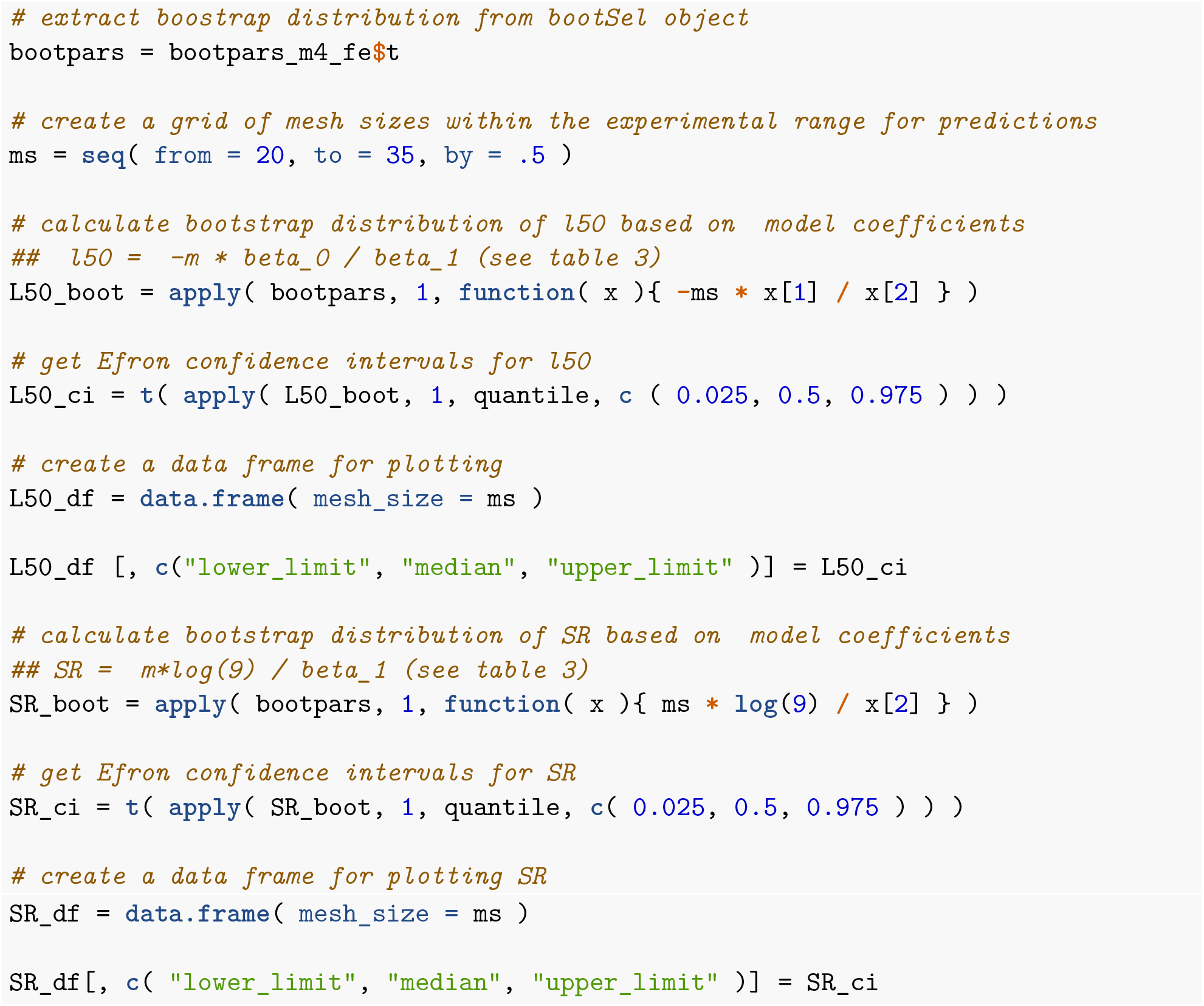

The following figure compares the predictions of *I*_50_ and *SR* estimated by model 4, with those from Santos et al. (2018). The black points in the figure represent values of *I*_50_ and *SR* estimated at haul level, and used in the original study as input data. Average predictions for *I*_50_ and estimation uncertainty (expressed in the amplitude of the confidence band) by model 4 are equivalent to those from the original study. Model 4 predicted higher values of *SR*, with larger uncertainty than Santos et al. (2018). This is a plausible result considering the different model structures applied, and the large variation of the by-haul estimates. Moreover, there is not statistical evidence to reject the possibility that the true value of *SR* could fall within the continue region of confidence bands overlap.

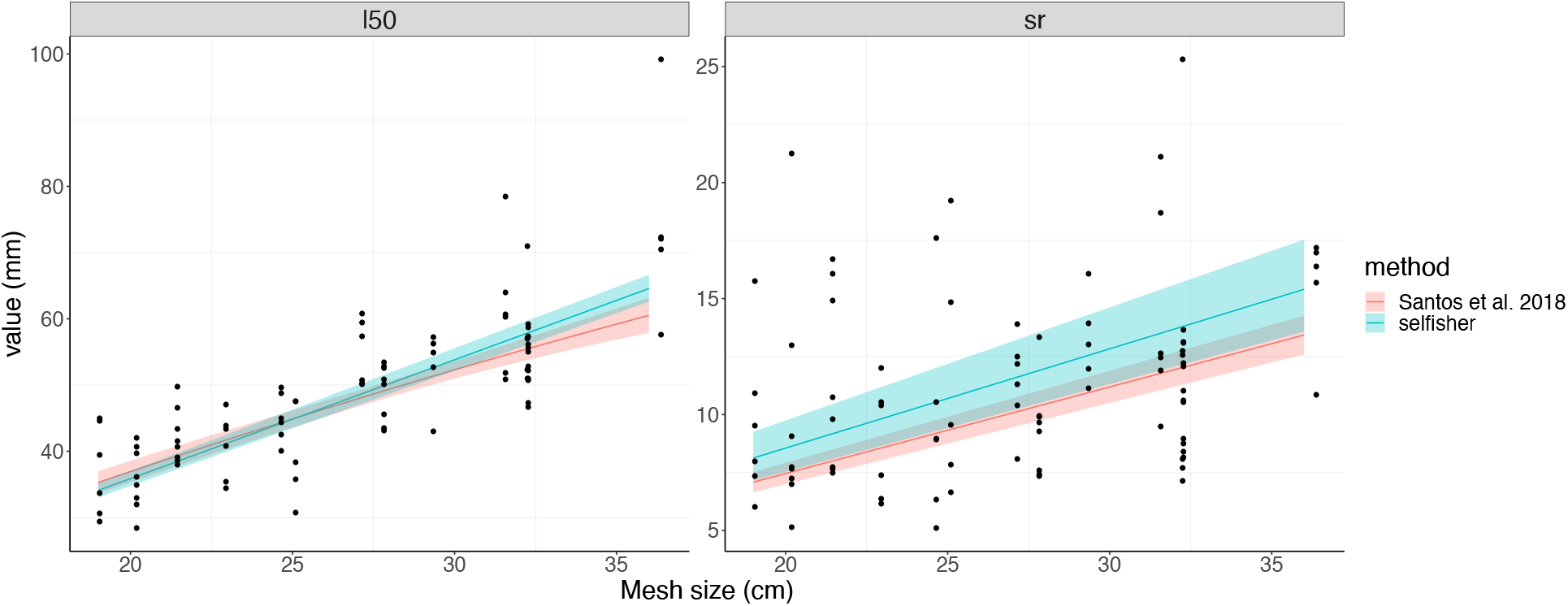

**Table 1:** Calculations to obtain selectivity parameters from population-average models 1-8 (fitted leaving out the random effects). Note that the last row for models 5-8 is the extension of model 4 to include additional covariates *x_i_, i* = 2, …, *p*.

| model | l50 | SR |
| --- | --- | --- |
| 1 | $\frac{-\alpha}{\beta_1}$ | $\frac{\log(9)}{\beta_1}$ |
| 2 | $\frac{-(\alpha + \beta_2 * ms)}{\beta_1}$ | $\frac{\log(9)}{\beta_1}$ |
| 3 | $\frac{-(\alpha + \beta_2 * ms)}{\beta_1 + \beta_3 * ms}$ | $\frac{\log(9)}{\beta_1 + \beta_3 * ms}$ |
| 4 | $\frac{-\alpha * ms}{\beta_1}$ | $\frac{\log(9) * ms}{\beta_1}$ |
| 5-8 | $\frac{-(\alpha + \sum_{i=2}^p \beta_i * x_i) * ms}{\beta_1}$ | $\frac{\log(9) * ms}{\beta_1}$ |

## Appendix 3 Catch comparison on unpaired hauls

This example deals with data from an experiment originally published by Savina et al. (2017). Two soak tactics, i.e., 12h at day and 12h at night, were compared in the Danish gillnet plaice fishery to estimate whether a change in soak tactics could help to catch less of the unwanted bycatch, i.e., the invertebrate edible crab (*Cancer pagurus*). The method developed by Herrmann et al. (2017) for assessing the relative length-dependent catch efficiency effect of changing from soak tactic *Day* to *Night* was used. This example is representative of experimental fishing where the catch data obtained for two different gear designs were not collected in pairs, and can allow for a different number of deployments.

### Preliminaries

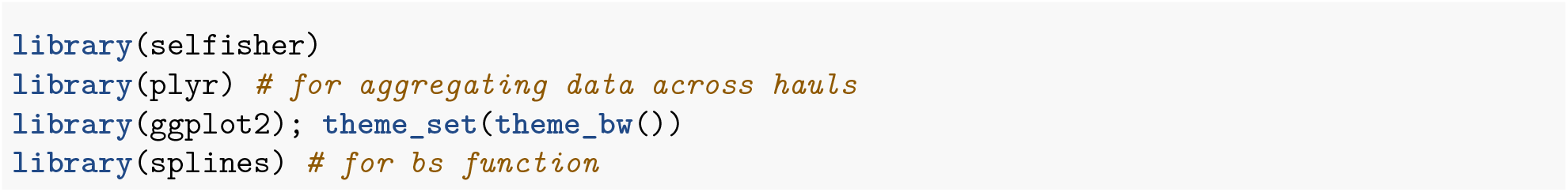

### Data structure

Load the data and check out the variables. This is a subset of the original dataset (one species, two soak durations). The data contains the length measurement of each individual to the nearest mm below (carapace width), as specified in the column “width”. Every day for 7 days (I to VII), three fleets (each consisting of three gillnets tied together, and labelled A, B and C) were soaked for 12 h during the day (*Day*) and three others during the night (*Night*). Each deployment of a fleet is considered as a “haul” (with haul name written as Day_Soak_Fleet). Gear unit design is the soak tactic, specified in the column “tactic”, with two levels: 12h at day (*Day*) and 12h at night (*Night*). “total” gives the number of individuals for each length class and haul. There was no sub-sampling.

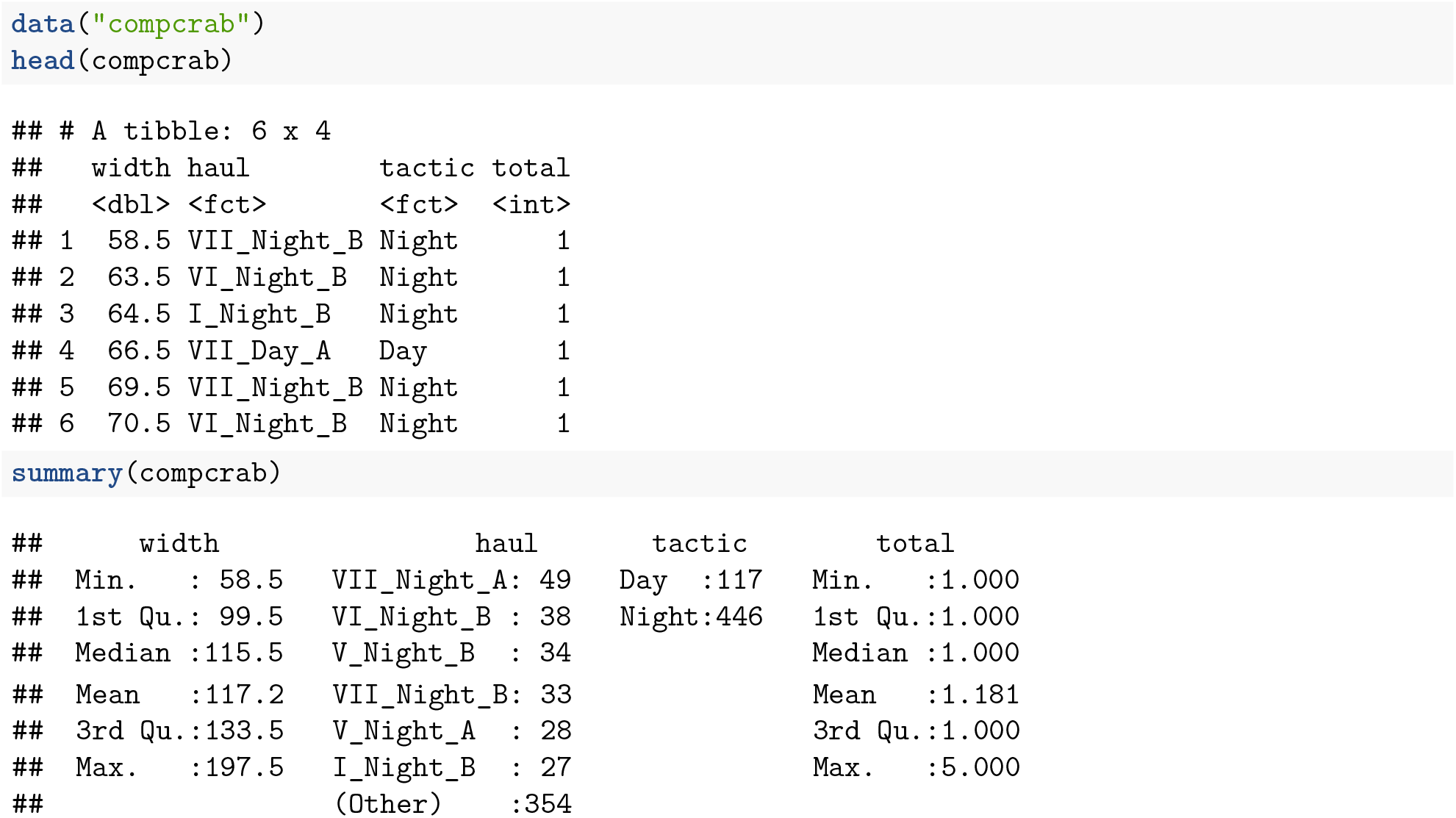

Here we can see that all hauls are contained in one data frame, organized into what is called “long format”, with *Day* and *Night* one after the other (unpaired).

### Transforming data

For a model in `selfisher`, we need to convert counts into proportions and totals. We use the ‘ddply’ function to calculate the proportion of fish entering one of the gear design (here *Night*) for each length class and haul, i.e., 1 for *Night* and 0 for *Day*.

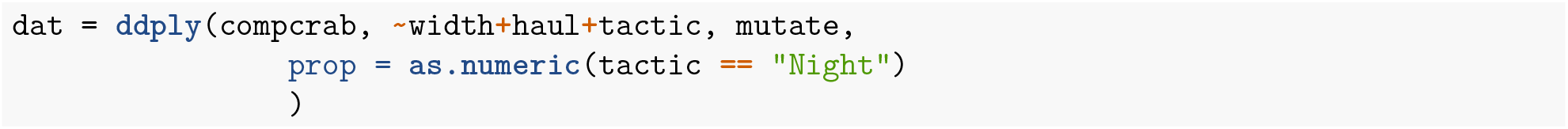

### Catch comparison

The following is a typical model for catch comparison of multiple haul data without subsampling using spline with the bs function.

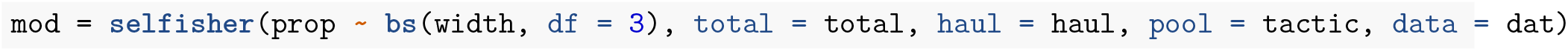

This models the proportion of fish in *Night* versus *Day* (prop) as a function of width. The `selfisher` function takes the total number of fish in *Day* and *Night* using a separate argument, total. The argument haul needs to be specified in order to perform double-bootstrapping as demonstrated below. Otherwise, it could be omited from the model specification as it doesn’t affect the fit. The haul argument tells the software how to group the data for resampling in the bootstrapping procedure. pool represents the different pools of hauls, i.e., one for each soak tactic, that is used in double bootstrap to produce same number of hauls by pool. Indeed, because the catch data obtained for *Day* and *Night* were not collected in pairs (and may not have the same total number of deployments), we sum data of the deployments carried out with *Day*, and data of the deployments carried out with *Night*.

Then we create a new data set to make predictions on.

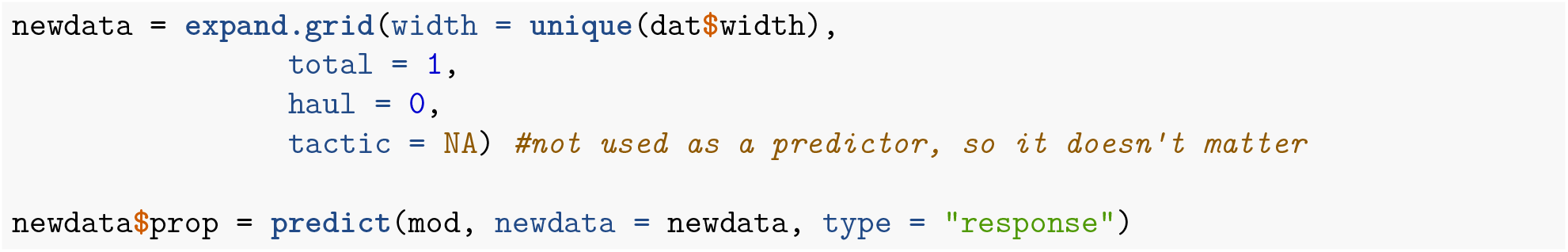

### Bootstrap to get CI on predictions

The code below runs in parallel on Mac and Linux computers as written here, but a Windows version was given above. This call to the function bootSel predicts the response variable based on the model mod and the covariates in newdata. Then we calculate the quantiles of the bootstrapped response variable, and transform the proportion into a catch ratio.

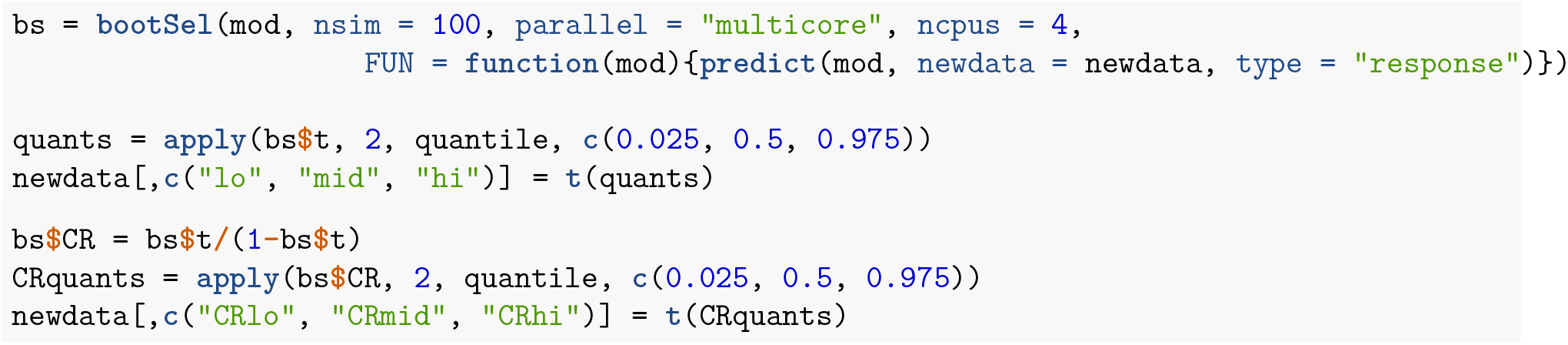

### Plot predictions

For plotting, we need to aggregate the hauls.

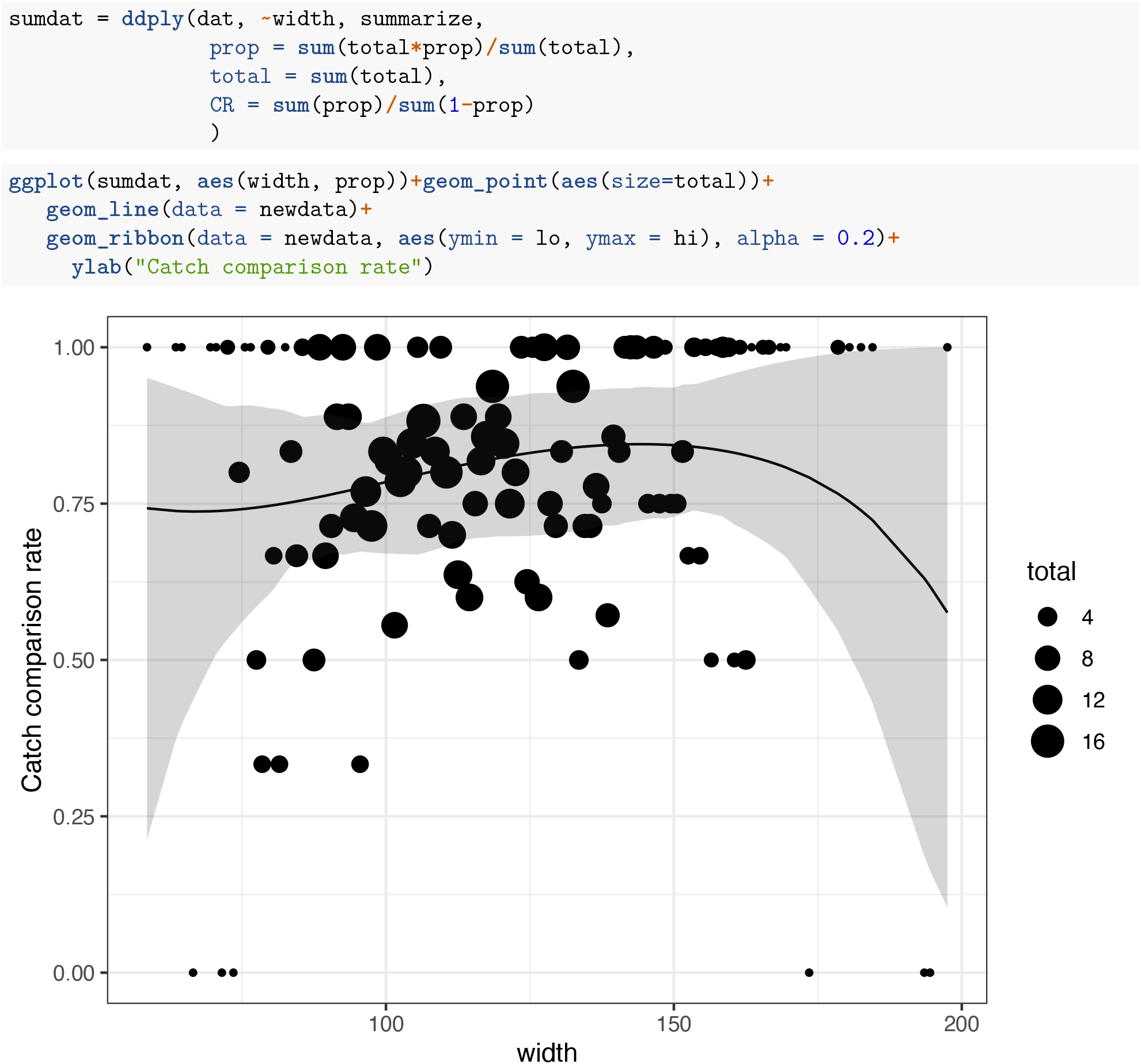

A graphical comparison to the published results in Savina et al. (2017; in red) shows that the estimated catch comparison curves and relative CIs are very similar.

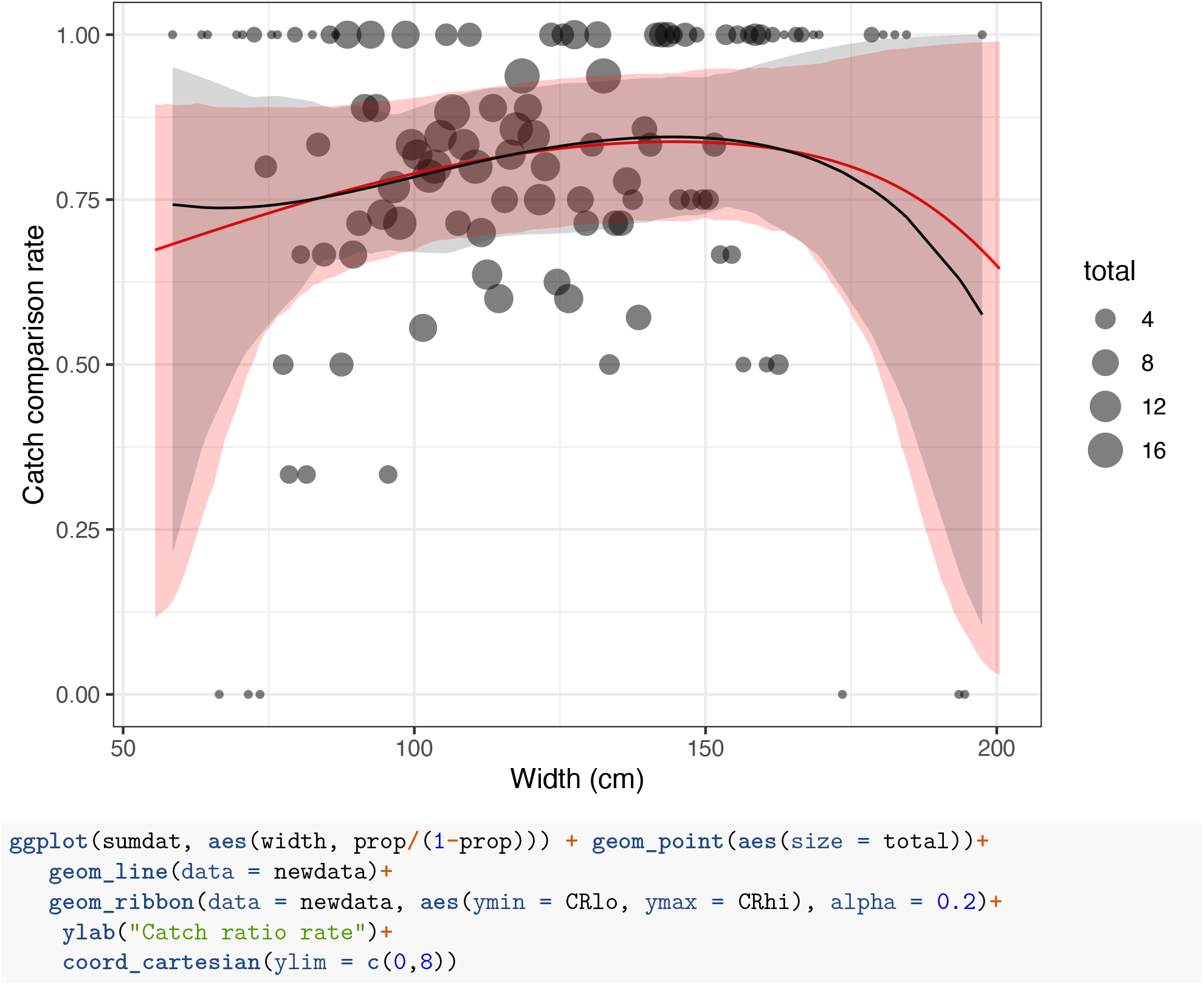
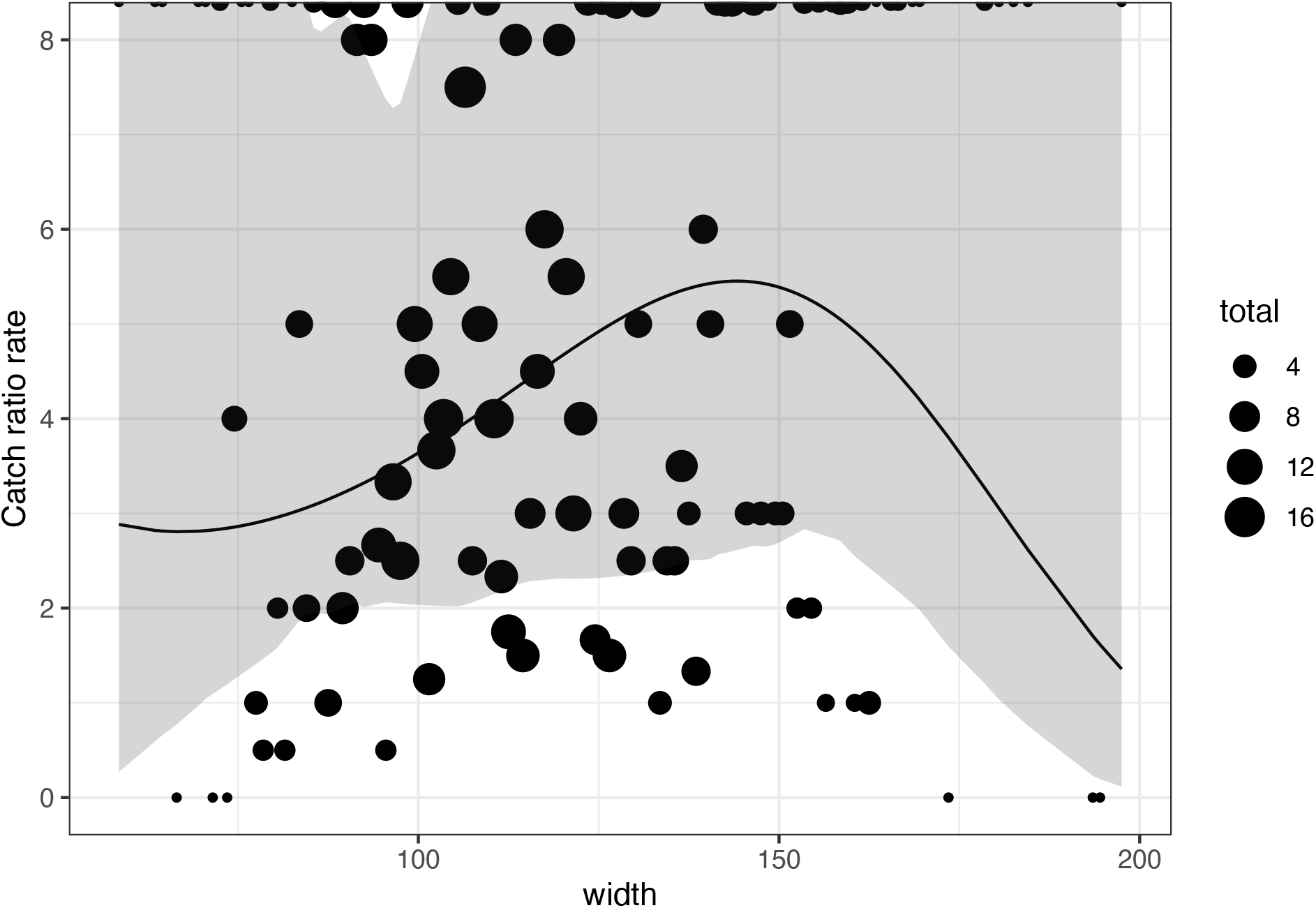

The catch comparison curves properly reflected the trend in the experimental points. The experimental rates were subject to increasing binomial noise outside the length classes representing the main bulk of the catches. The results for edible crab showed significantly higher catches for 12 h at night compared to 12 h at day. On average, there were four times more catches for 12 h at night than 12 h at day. There was no strong indication of a length dependency in the data.

## Appendix 4 Catch comparison analyses of paired hauls of *Nephrops* twin-rigged trawls

This example is based on the data from Melli et al. (2018). An anterior gear modification, namely the counter-herding device FLEXSELECT, was tested in a twin-rig configuration, where two identical bottom trawls were towed in parallel. One trawl was equipped with FLEXSELECT, while the other worked as baseline. The aim of the study was to determine if FLEXSELECT could prevent fish species from entering the trawl in a *Nephrops (Nephrops norvegicus)* fishery. Data for haddock *Melanogrammus aeglefinus* were collected for 21 hauls. Of these, 13 were conducted in day-time and 8 in night-time. In each haul and for each trawl, the total length (rounded down to the lower centimitre) of all haddock individuals was recorded.

### Preliminaries

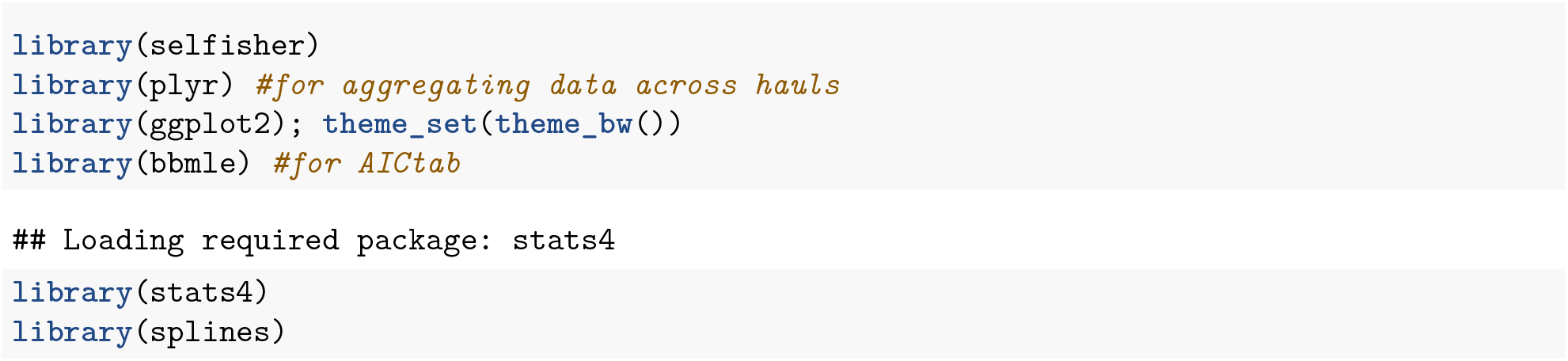

### Data structure

First, we load the data and check out the variables. Here we can see that all the hauls are contained in one data frame (long format), with each row corresponding to a length class in a given haul. The number of individuals of that length-class caught in each trawl is reported in the columns `TEST1` (test trawl) and `TEST2` (baseline trawl). The column `TIME` classifies the haul as day-time `(D)` or night-time `(N)`.

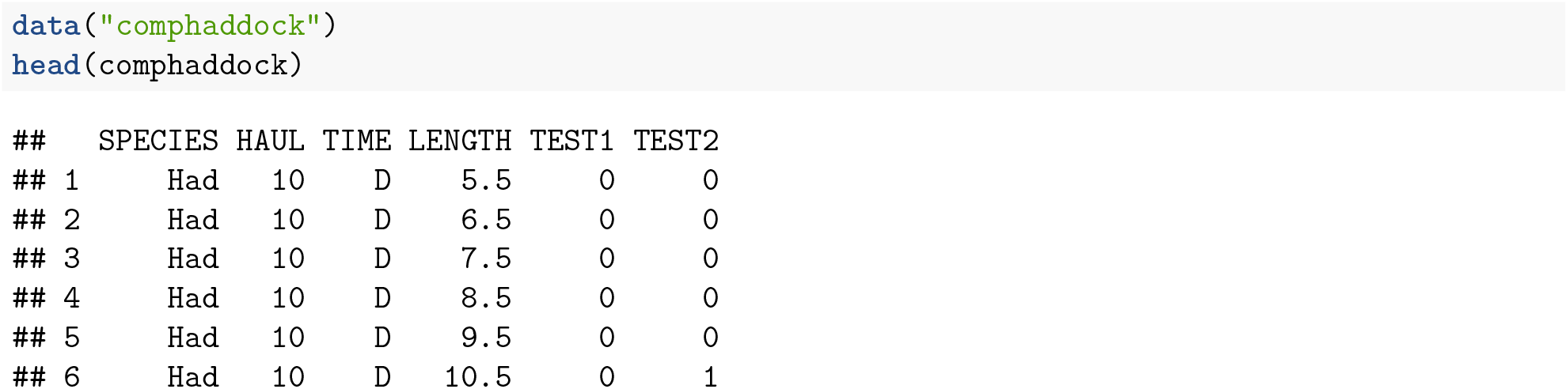

To understand if the test trawl caught significantly fewer individuals of a given length-class than the baseline trawl we need to perform a catch comparison analysis (Krag et al., 2015). This analysis estimates the probability of catching an individual of a given length in the test trawl given that it was caught in either trawl.

In addition, the analysis aims at determining if the length-based efficiency of FLEXSELECT presents diel differences, as haddock is known to migrate vertically in the water column during the night.

### Transforming data

Before fitting a model in `selfisher`, the following preparatory steps need to be performed:

1. Convert counts (i.e. number of individuals caught per length-class) to totals, proportions and ratios; This step is required because, unlike with other GLM functions for binomial regression, it is not possible in `selfisher` to specify the binomial variable as a two-column response variable, e.g. cbind(`N_TEST1, N_TEST2`).
2. Remove eventual length classes where no individuals were caught;
3. Scale the length. This step is necessary for numerical stability, as a model often used for catch comparison analyses is the polynomial of order four, which requires to raise the length to the 4th power.

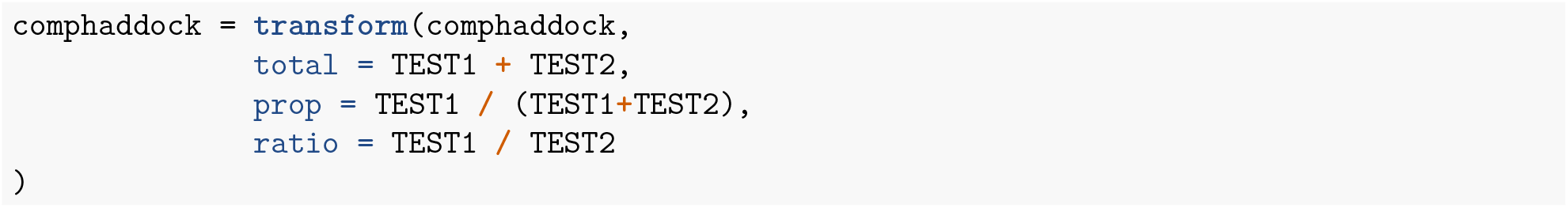

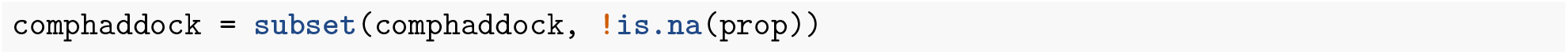

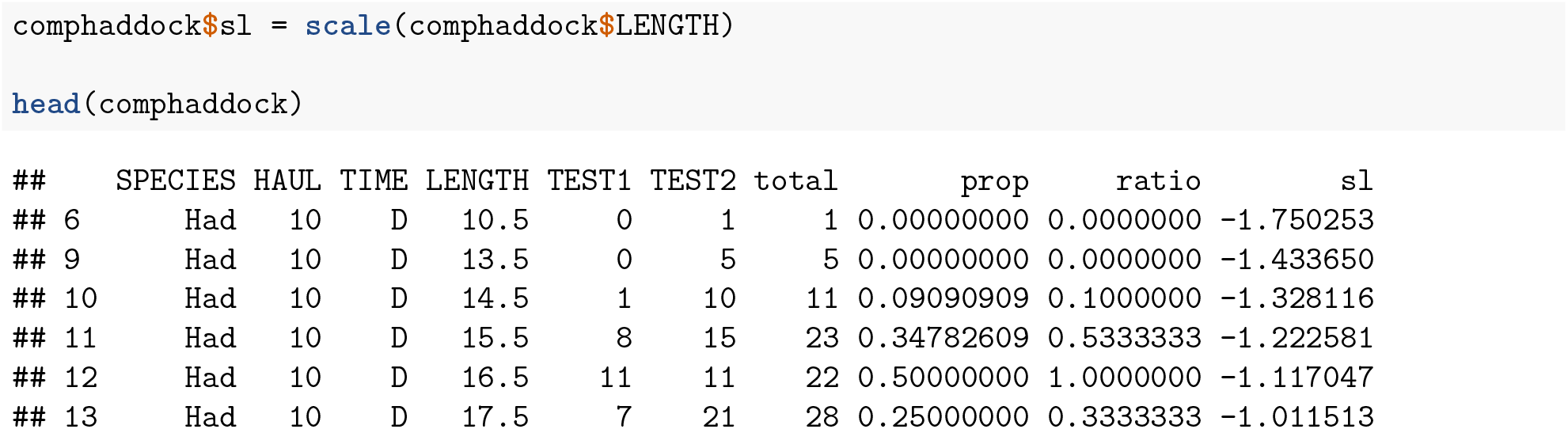

### Model fitting

The following is a typical model for catch comparison data with multiple paired hauls, which models the proportion of fish in the test versus the baseline trawl (`prop`) as a function of length (sl). This is epressed in the `selfisher` function by a two sided formula with the proportion (`prop`) on the left side and fixed and random effects on the right side. Formulas in the `selfisher` package follow the convention of the lme4 and `glmmTMB` packages.

Since we are interested in determining if there is a length-dependent difference in the efficiency of the Test gear between day-time and night-time, we include in the model `TIME` as an explanatory variable.

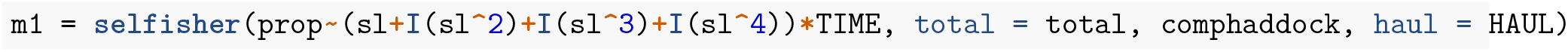

In this example all individuals were length-measured (i.e. there was no subsampling). In case of a subsampled species, an offset or q-ratio needs to be specified in the model. The `selfisher` function takes the total number of fish in the test and baseline using a separate argument, total. The argument haul needs to be specified in order to perform double-bootstrapping as demonstrated below. Otherwise, it could be omited from the model specificaiton as it doesn’t affect the fit. The haul argument tells the software how to group the data for resampling in the bootstrapping procedure.

### Alternative model

An alternative approach would consist in fitting a spline (Miller, 2013), often preferred to polynomial interpolation because it yields similar results while avoiding Runge’s phenomenon (i.e. oscillation at the edges of the length range represented in the data).

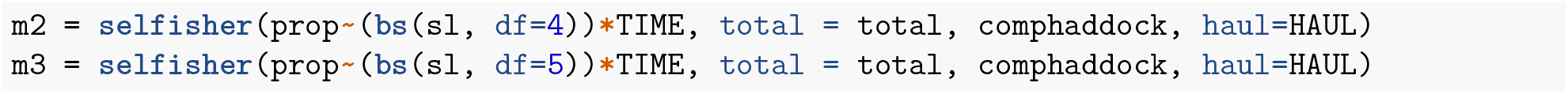

### Model comparison

We can determine which model fits best using the Akaike’s Information Criterion (AIC; Akaike, 1974).

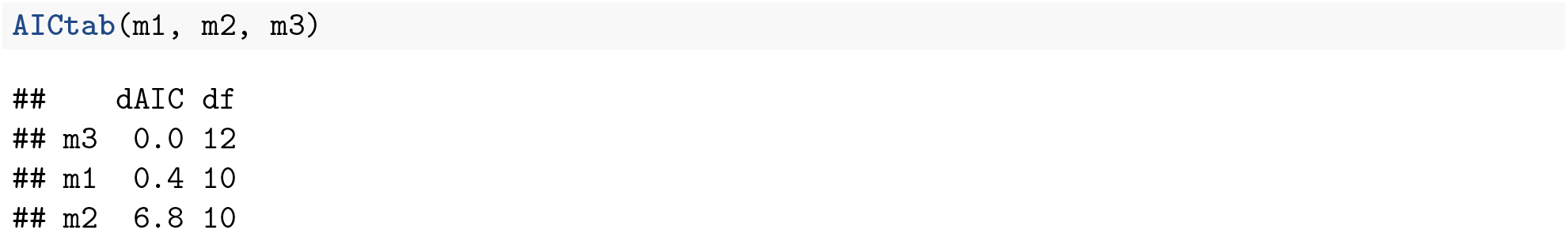

This tells us that m1 and m3 show equally good fit (0.4 delta AIC units). According to the parsimony rule, we selected m1 that is a simpler model.

### Predictions

To see how the model fits the data, we need to plot observations and predictions together, keeping them separated by TIME.

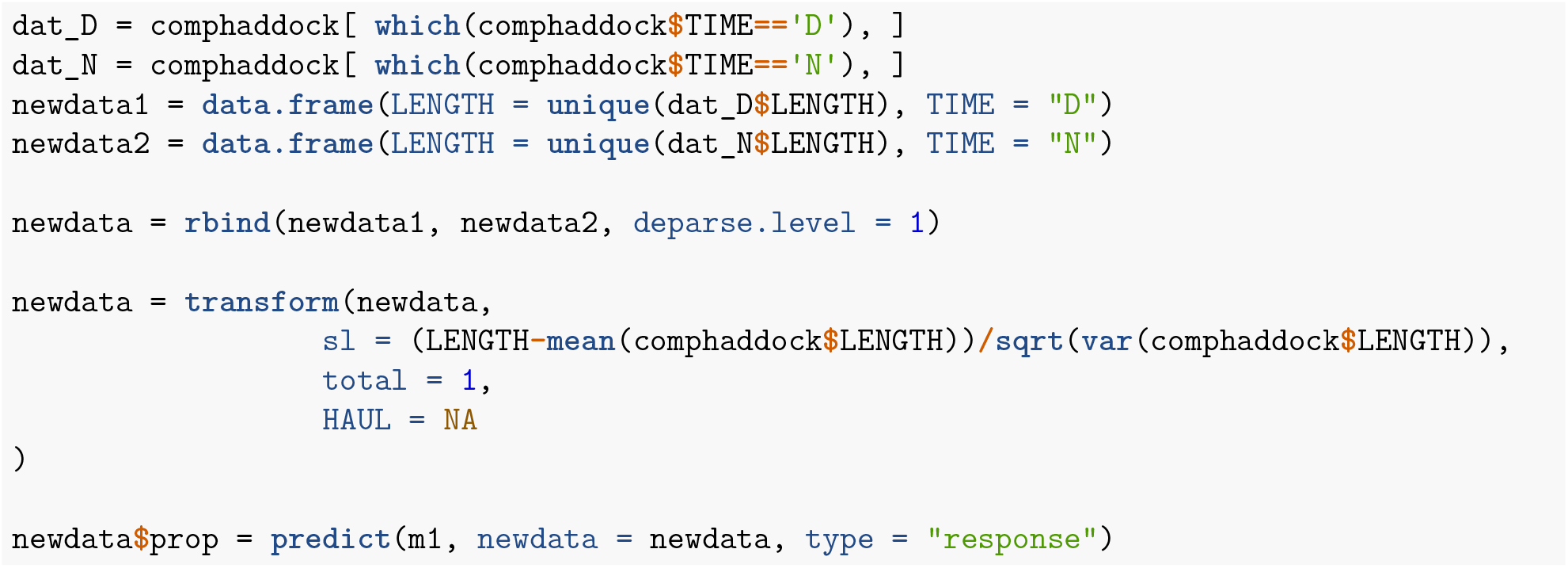

### Confidence intervals by double-bootstrapping

We then estimate the 95% Efron Confidence intervals (Efron, 1982), by accounting for within- and between-hauls variation (Millar, 1993). The code below resamples hauls, then resamples fish within hauls, fits the model to the resampled data, then makes predictions from the model onto `newdata`. The type of predictions we want in this case are the catch comparison rates, thus we specify type=“response”. To read about the predict function, `type ?predict.selfisher` in the R console.

### Windows bootstrapping in parallel

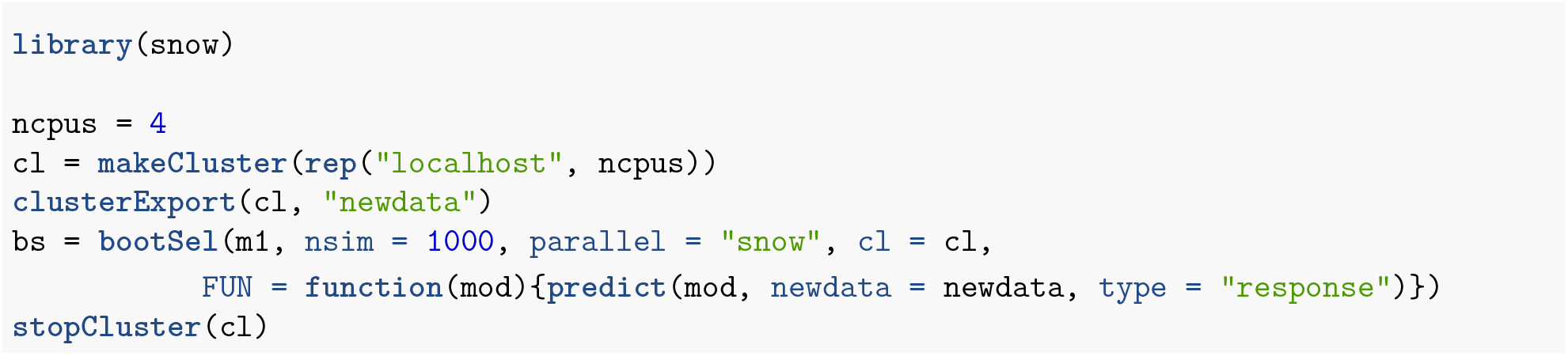

Code for bootstrapping in Mac and Linux

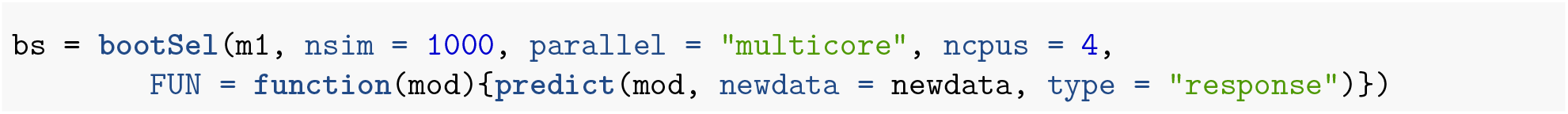

Then we calculate quantiles across bootstraps for each row of newdata.

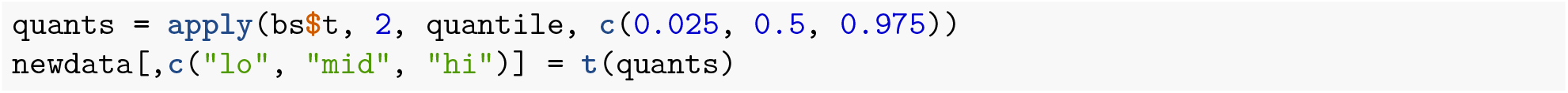

### Plotting with CIs

Here, we plot the modelled catch comparison curve with CIs and experimental obsevations, obtained by aggregating the hauls per TIME.

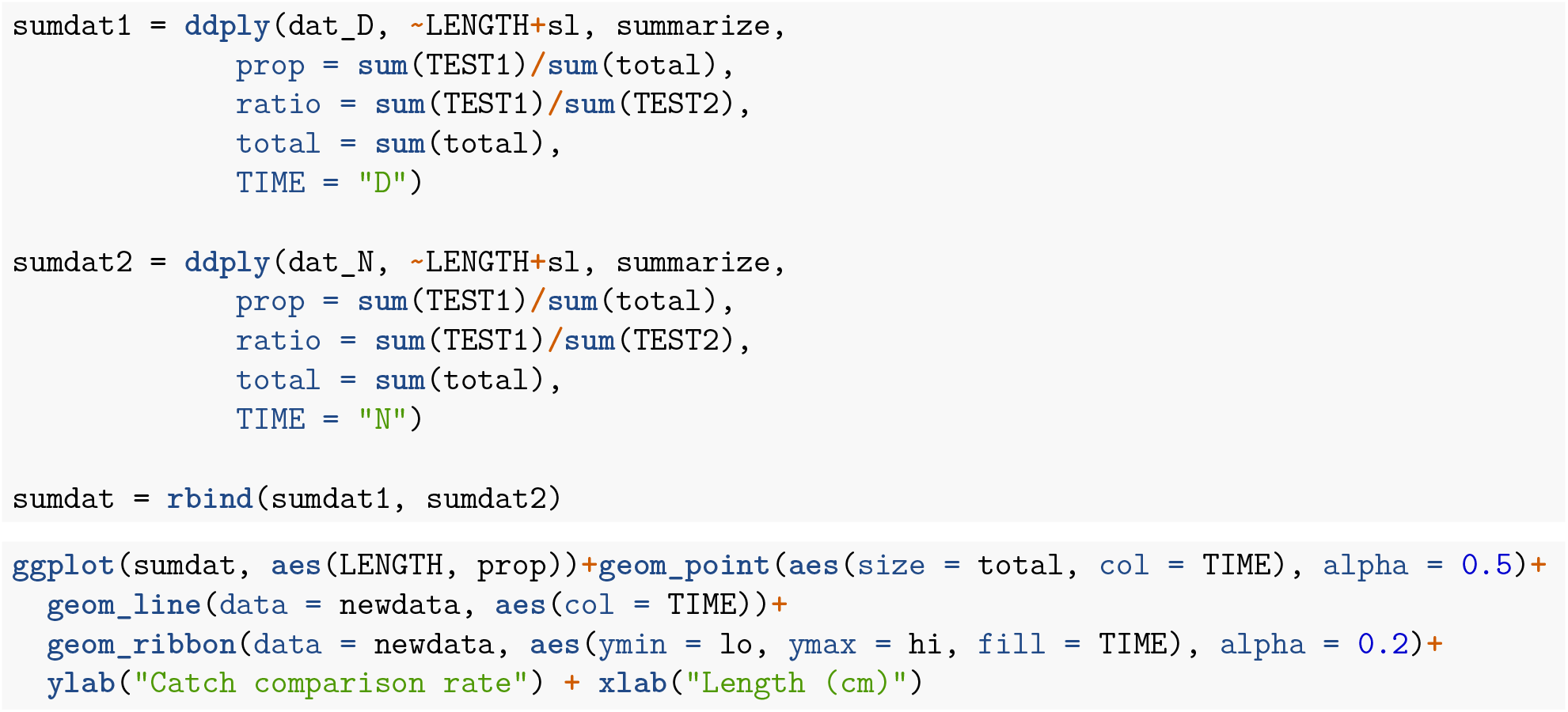
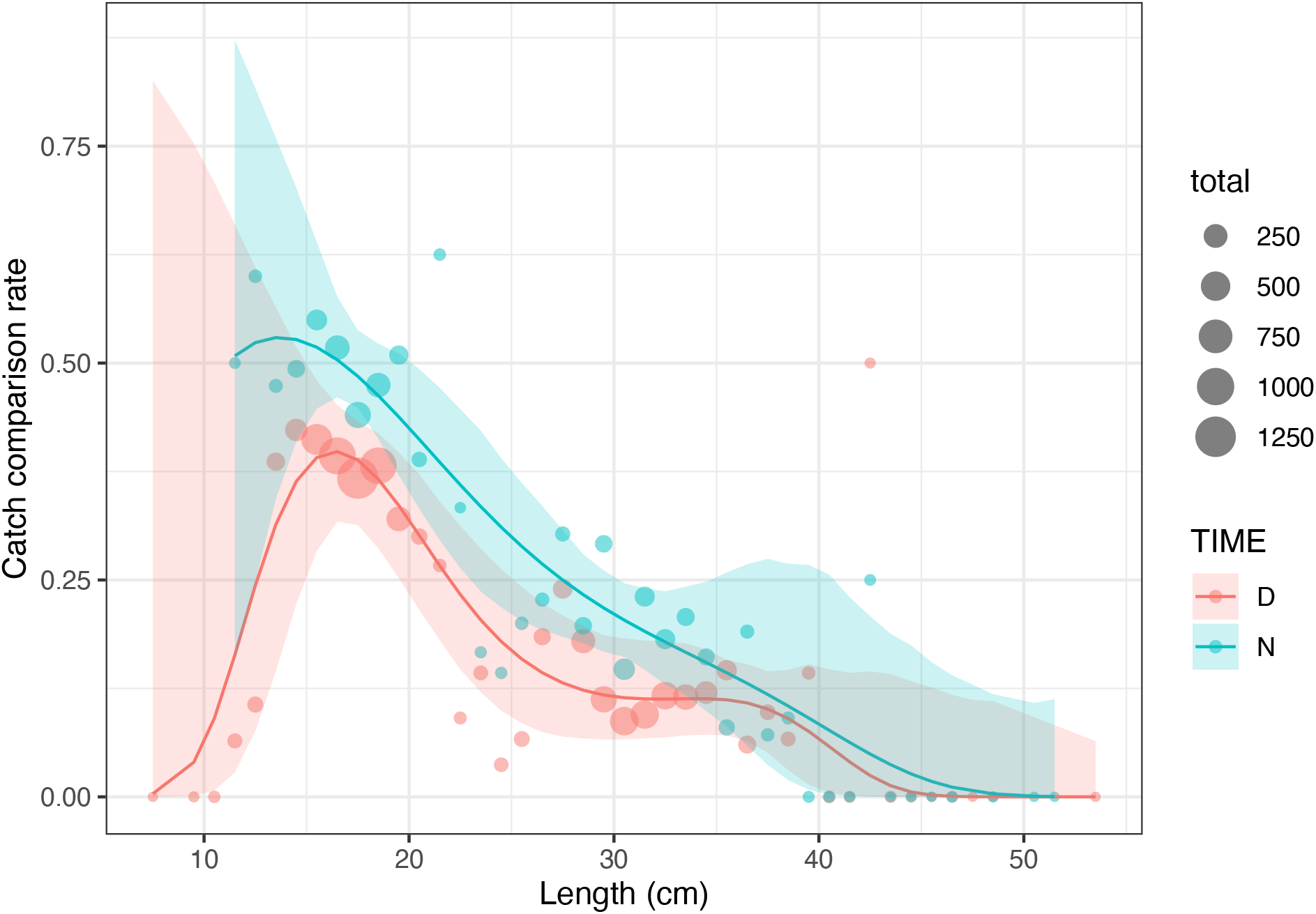

In accordance to Melli et al. (2018) a significant difference in catch comparison rate between day-time and night-time is found for individuals between 16 and 18 cm, as represented by the lack of overlapping between the CIs.

Melli et al. (2018) argued that, being the difference in a length range that is not usually retained when using a commercial codend, and that commercial fishing operations take place in both day- and night-time conditions, it is of greater interest to estimate the effect of FLEXSELECT with respect to the baseline trawl without the factor TIME.

Therefore, we repeat the steps of the process leaving out the factor TIME.

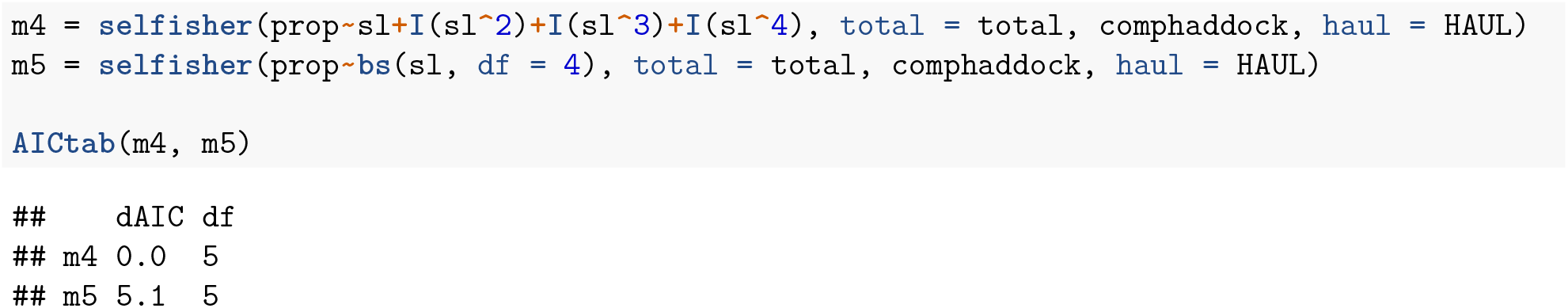

Again, we select the polynomial of order 4 and use it to predict the catch comparison rates with CIs, which are then plotted against the overall experimental observations.

### Windows code

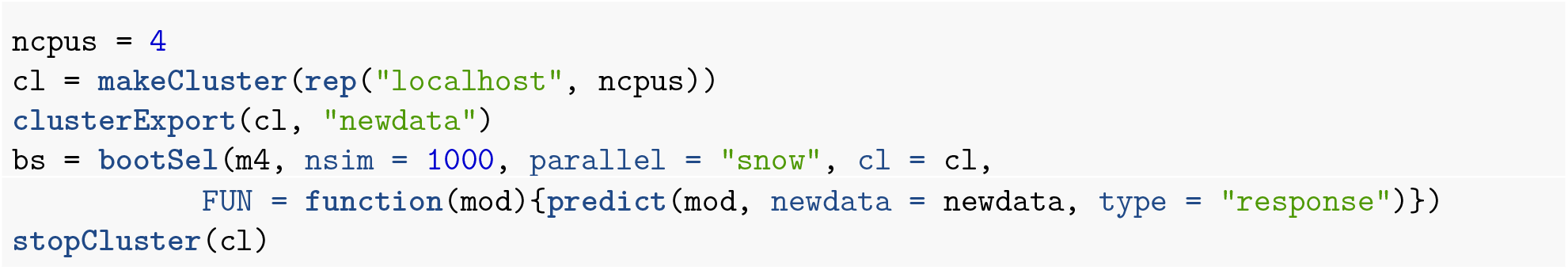

### Mac and Linux code

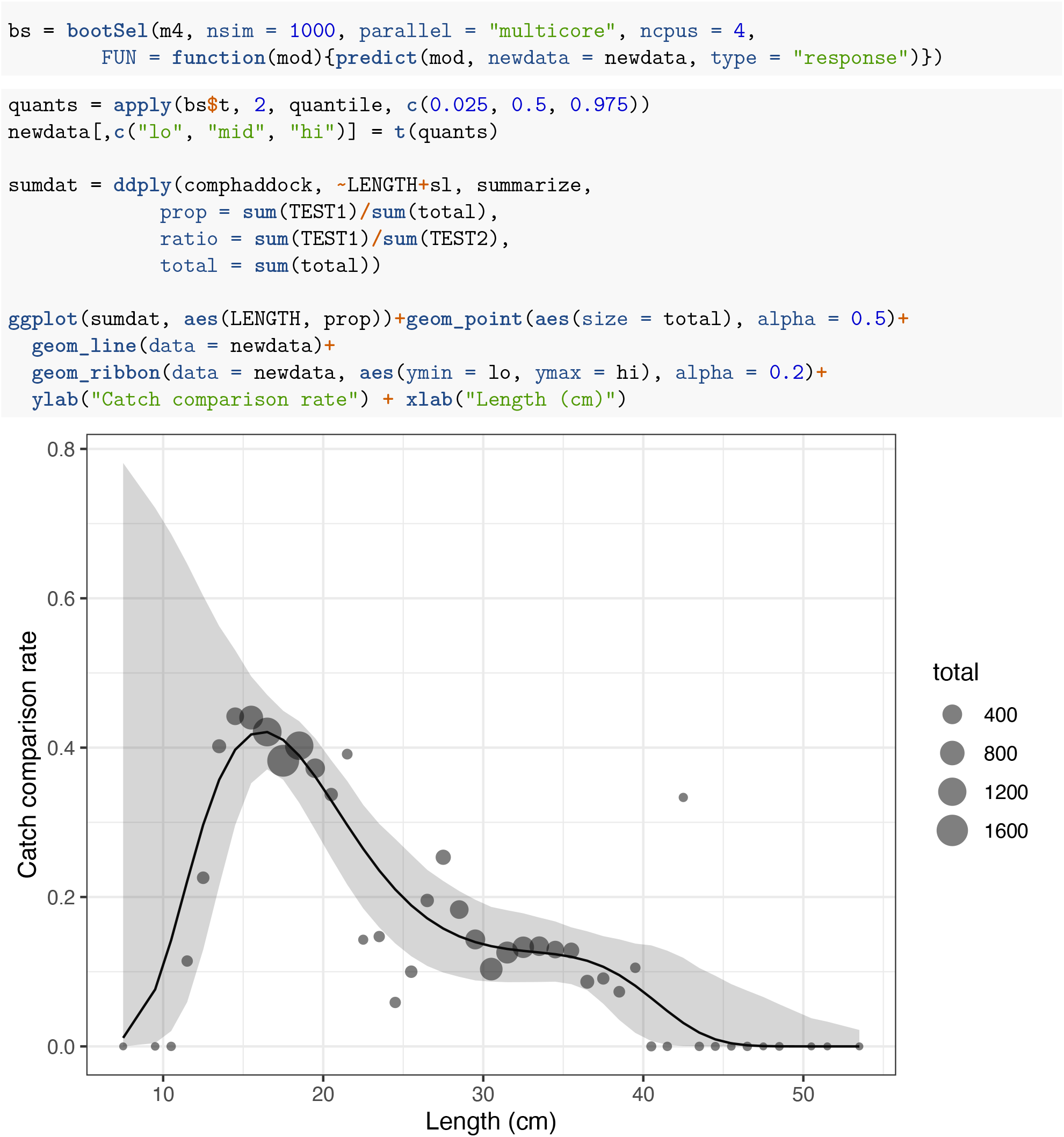

A graphical comparison to the published results in Melli et al. (2018; in red) shows that the estimated catch comparison curves and relative CIs are very similar.

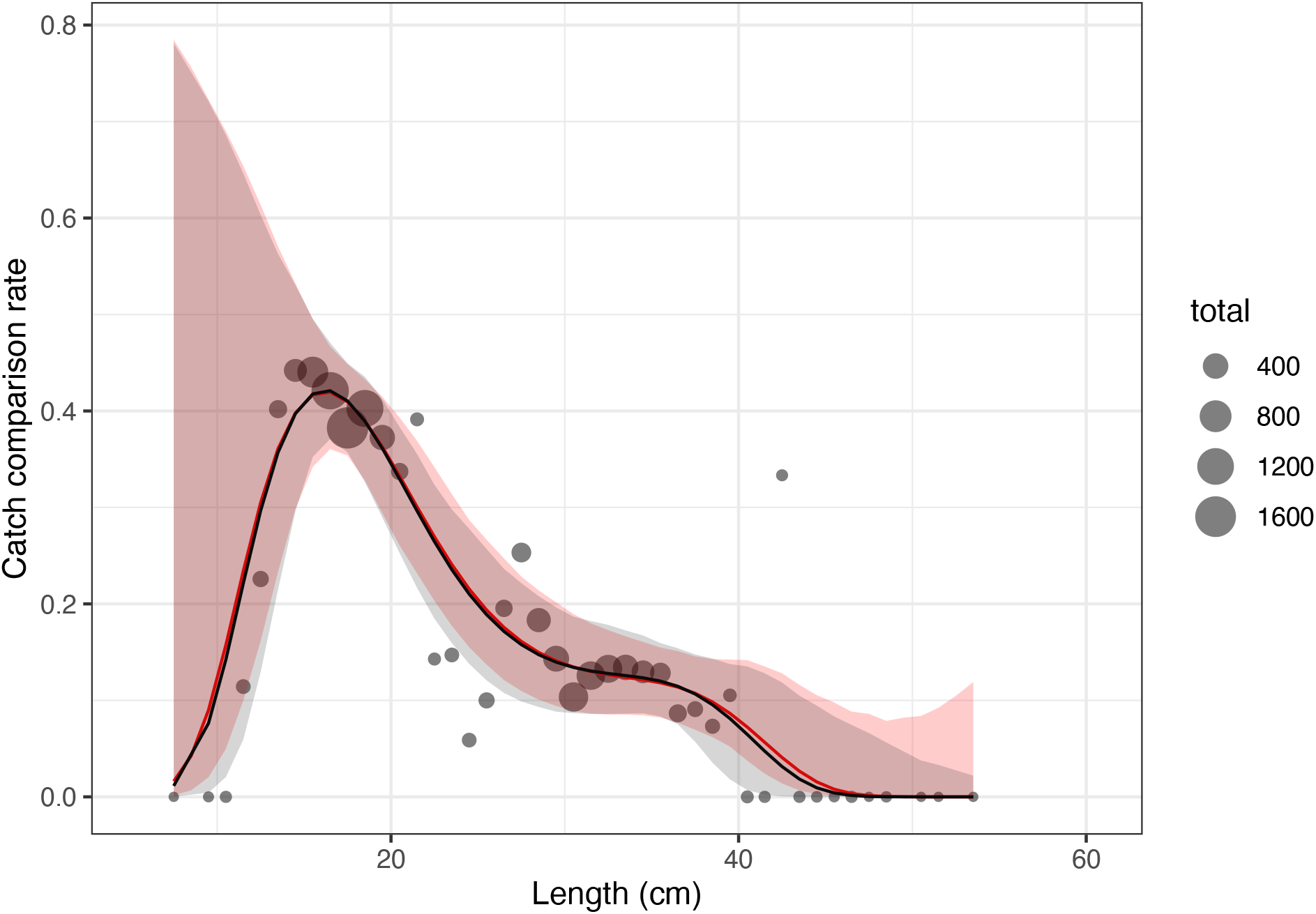

### Catch ratio

To directly quantify the difference in catch between the test and baseline trawls, it is common practice to estimate the catch ratio curve, using the relationship between catch comparison rate (cc) and catch ratio (cr): cr=cc/(1-cc)

First, we obtain predictions for the catch ratio specifying `type=“ratio”` in the predict function.

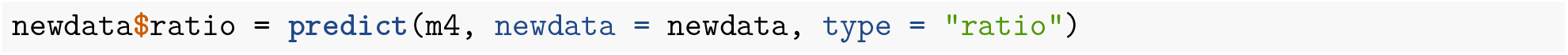

Second, we apply the relationship between cc and cr to obtain the CIs for the catch ratio curve.

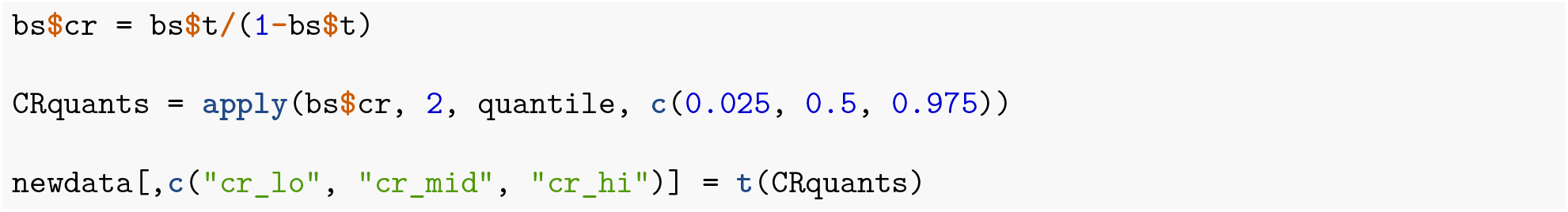

### Catch ratio plot

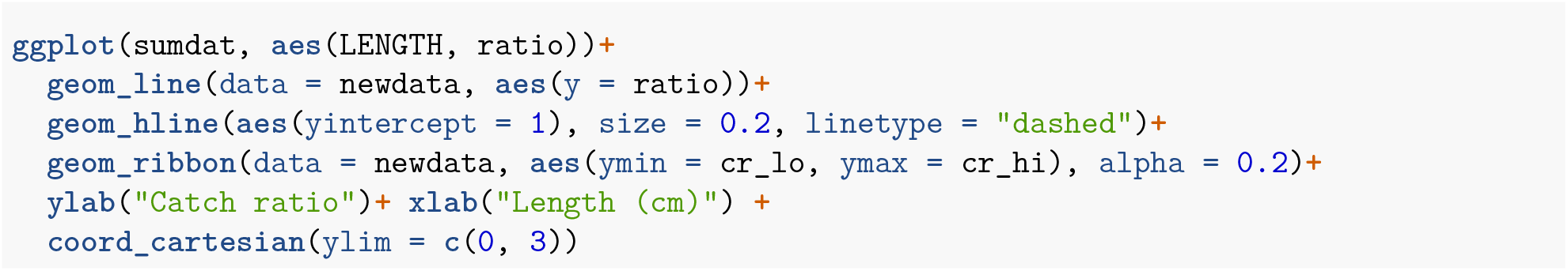
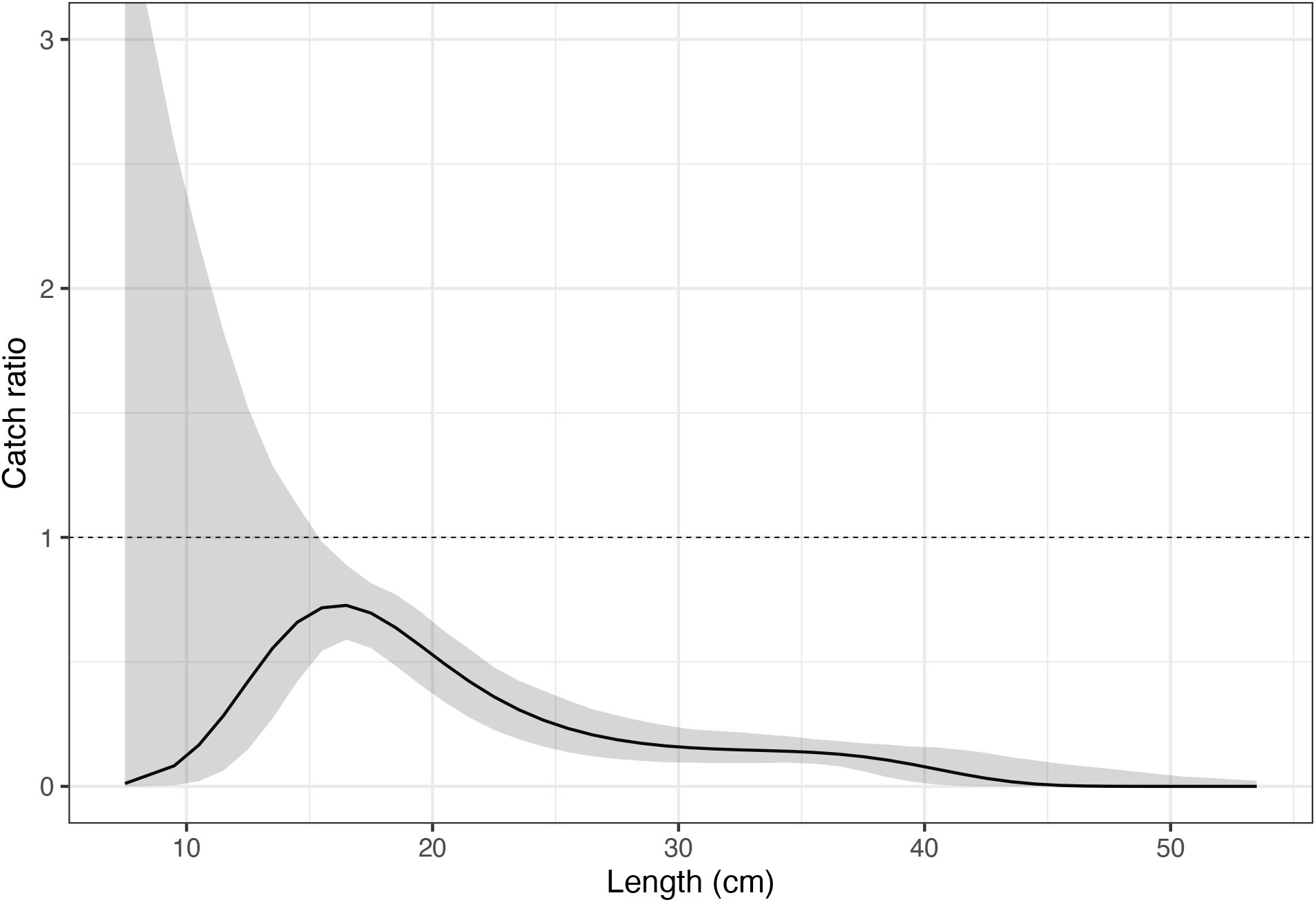

The results show that for individuals above 16 and up to 53 cm, the test gear with FLEXSELECT retained significantly fewer individuals. The effect is length-dependent, with a more pronounced reduction at larger length classes. Considering a minimum conservation reference size of 27cm for haddock in the fishing area (Skagerrak and Kattegat), the reduction of commercial-sized individuals is above 60%.

